# Physiologically Informed PCA-Partial Correlation for highly Collinear Brainstem fMRI Networks

**DOI:** 10.64898/2026.06.08.730883

**Authors:** Santa Sozzi, Alejandro Luis Callara, Simone Cauzzo, Enzo Pasquale Scilingo, Paola Binda, Nicola Vanello

## Abstract

Functional connectivity (FC) approaches from resting-state fMRI (rs-fMRI) are amply spread to investigate the cortical organization, yet the brainstem remains relatively underexplored despite its pivotal roles in both physiological and pathological conditions. The highly collinear network, in which the strongly interconnected nodes and the widespread neuromodulatory influences induce indirect or mediated interactions, make the estimation of direct brainstem FC challenging. Standard bivariate methods fail to recover the true network structure in such complex topologies, causing false positive interactions. On the other hand, partial correlation can potentially estimate the direct FC, but multicollinearity issues and collider-induced spurious correlations limit its application in high-dimensional scenarios. Here, we propose a physiologically informed framework in which the conditioning strategy for partial correlation estimation is tailored for the investigation of the brainstem and its direct interactions within the network and with whole-brain regions. Specifically, we employed a PCA-regularized partial correlation (PCA − ρ_PC_) approach, where PCA is applied to the brainstem covariates to mitigate multicollinearity and model shared modulatory variance. We show that PCA − ρ_PC_ improves the robustness and interpretability of brainstem FC, yielding sparser and more physiologically plausible connectomes compared with conventional (regularized) approaches. Both simulation and real fMRI data raise the possibility that Pearson’s and PCA-regularized approaches may complement each other in an effort to unravel the pattern of direct vs. indirect effects in highly collinear settings, paving the way for future extensions in a wide range of multivariate neuroimaging applications.

## Introduction

Resting state functional connectivity (rs-FC) analysis is widely used to investigate large scale brain organization from fMRI data. Statistical dependencies between regional BOLD timeseries are extensively employed to characterise intrinsic functional networks, inter-individual and longitudinal variability, and disease related alterations in brain dynamics [1], [2], [3]. Extensive methodological works on quality control and processing pipelines have been carried out to reliably estimate cortical FC [4], [5], [6], [7], but brainstem FC is still relatively underexplored. This gap is startling given its physiological and clinical relevance: the brainstem plays a pivotal role in autonomic regulation, arousal, sensory processing, motor coordination, and neuromodulation [8], [9], [10], as well as a potential driver or modulator of disease progression in neurodegenerative and neuropsychiatric conditions [11], [12], [13]. Despite this broad impact, mapping brainstem FC from rs-fMRI is challenging. Conceptually, the brainstem poses a demanding statistical problem for modeling functional interactions. Methodologically, its networks are characterized by widespread connectivity with cortical and subcortical regions [14] and by a highly interconnected internal organization. Moreover, brainstem nuclei exert strong and often overlapping neuromodulatory influences, resulting in a system with strong collinearity and complex interaction motifs, as already observed in the attempts at delineating the brainstem-to-cortex functional connectome [15], [16]. In this context, individual nuclei may concurrently contribute to shared variance across multiple regions (confounders), mediate indirect activity between non-directly interacting areas (element of causal chain systems), or integrate convergent inputs arising from different functional systems (collider). Therefore, brainstem-to-brain and brainstem-to-brainstem FC patterns do not only reflect direct physiological interactions but may also emerge from indirect pathways mediated by other regions or from shared modulatory processes.

The most commonly used FC metric, Pearson correlation (*ρ*_*P*_), characterizes overall (mediated) statistical dependence but cannot isolate direct from indirect interactions. Shared inputs, multi-step chains, and global modulatory fluctuations can inflate pairwise correlations, resulting in dense and potentially redundant connectomes [17]. Partial correlation (*ρ*_*PC*_) provides a multivariate alternative by estimating the association between two regions while taking into account (conditioning) the activity of the remaining network regions [18], [19], becoming a more plausible candidate for “direct” FC. However, its estimation can be inflated by multicollinearity (i.e., strong mutual correlations among the predictor variables) [20]. In highly collinear systems, both regression-based and precision matrix-based *ρ*_*PC*_ estimation becomes ill-conditioned, leading to unstable parameters with inflated variance and sensitivity to noise [20], [21]. Moreover, conditioning on collider nodes can induce artificial dependencies between otherwise independent variables [22], [23], [24], leading to spurious conditional associations [17], [23] (Berkson’s paradox). Several regularization strategies have been proposed to improve the robustness of FC estimation through *ρ*_*PC*_. These include sparse inverse covariance estimation methods such as graphical Lasso, graphical Ridge, and Elastic Net regularization [25], as well as alternative formulations such as minimum partial correlation [23]. More recently, Peterson et al. compared pairwise correlation, partial correlation, graphical Lasso, graphical Ridge, and PC regression, demonstrating that regularized *ρ*_*PC*_ with cross-validated parameters significantly improves the reliability of FC estimates [26]. However, these regularization approaches have typically been developed and tested at the whole-brain level, where all regions are treated equivalently within the conditioning set [25], [27], [28]. Moreover, they are primarily driven by statistical optimality (e.g., sparsity, reliability) rather than by explicit modelling of how a specific set of nodes (e.g., brainstem nuclei) act as confounders or mediators in the network. In highly interconnected subcortical systems such as the brainstem, whole-brain conditioning increases the dimensionality and collinearity of the regression problem and may reduce interpretability by removing physiologically meaningful shared variance, thereby obscuring the contribution of brainstem-specific modulatory processes to large-scale functional interactions.

In this work, we propose a pipeline conceptually situated within regularized partial correlation but tailored to the brainstem and informed by domain knowledge of its neuromodulatory function. Specifically, our approach is based on a physiologically informed PCA-regularized partial correlation framework (*PCA* − *ρ*_*PC*_) [29] designed for highly collinear brainstem rs-fMRI networks and implemented in a principal component regression (PC-regression) logic. Given the high inter-connectivity among nuclei, their coordinated neuromodulatory dynamics, and their widespread influence on cortical and subcortical activity, rather than conditioning on the entire set of cortical and subcortical regions, our framework restricts the conditioning space to brainstem covariates, motivated by the hypothesis that a substantial portion of indirect brainstem-to-brain connectivity arises from shared brainstem modulatory activity. Principal component analysis (PCA) is then applied to the brainstem conditioning set to transform highly correlated signals into a low-rank orthogonal representation of the regressors. This strategy mitigates multicollinearity, improves conditioning of the regression problem, and models dominant shared variance within the brainstem network. To further reduce spurious conditional dependencies caused by the collider effect, we elaborated from [17], [30] and jointly uses *PCA* − *ρ*_*PC*_ and standard pairwise correlation. The resulting framework aims to improve the robustness, interpretability, and stability of conditional FC estimation while preserving nucleus-specific interactions. It was evaluated on both simulated and experimental rs-fMRI data. Simulations included synthetic multivariate networks with confounders, colliders, and causal chains under increasing dimensionality and collinearity conditions. Experimental validation was performed using 3T resting-state fMRI data and probabilistic atlases of human brainstem nuclei [31], [32], [33]. We characterized *PCA* − *ρ*_*PC*_ performance against *ρ*_*P*_, standard *ρ*_*PC*_, and penalized regression approaches in terms of network recovery, conditioning, and runs stability. Our results show that *PCA* − *ρ*_*PC*_ improves the stability and interpretability of brainstem FC estimation by mitigating multicollinearity and reducing indirect or mediated interactions. These findings support the use of physiologically informed PCA-regularized partial correlation strategies for the investigation of highly collinear brain networks.

## Methods

In both simulation and experimental datasets, we developed a region-of-interest (ROI)-based framework to characterize brainstem FC and differentiate direct from mediated interactions. ROIs timeseries were arranged into a matrix, *X* ∈ ℝ ×(*B*+*CTX*), where *T* is the number of timepoints, *B* is the number of brainstem nodes, and *CTX* denotes the number of target cortical ROIs. In simulations we compared three connectivity estimators, *ρ*_*P*_, *ρ*_*PC*_, and *PCA* − *ρ*_*PC*_ (with and without collider correction), in both *ad hoc* toy examples and high dimensional synthetic networks to assess their ability to recover the ground truth (GT) connectivity structure in terms of true positive rate (TPR), false positive rate (FPR), false negative rate (FNR), and balanced accuracy (BA), while systematically varying network dimensionality, number of time points, noise level, and the dimensionality of the PCA-based conditioning set. Classical regularization approaches were not included, as these methods have already been extensively evaluated in previous literature on simulated brain networks and connectivity inference [31], [33], [35], [49]. In contrast, relatively few studies have investigated PC-regression schemes for FC estimation under controlled simulation settings [34] [50]. Experimental analyses additionally included regularized *ρ*_*PC*_ approaches based on Lasso (*ρ*_*L*1_), Ridge (*ρ*_*L*2_), and Elastic Net (*ρ*_*EN*_) penalties. These estimators were evaluated in terms of robustness to collinearity and their ability to yield physiologically plausible cortical and subcortical connectomes.

### A. Pearson and Partial Correlation

As a baseline FC measure, we used pairwise Pearson correlation coefficients computed as:

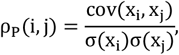

where x_i_ and x_j_ are the timeseries of ROIs *i* and *j*, respectively. Consequently, for each subject, we obtained a symmetric correlation matrix ρ_P_ ∈ ℝ (B+CTX) × (B+CTX).

To estimate direct associations while controlling for the influence of other signals, we computed naïve partial correlation via multiple linear regression (MLR). In our implementation, the covariate (predictor) matrix was defined using brainstem-only regressors, Z ∈ ℝ^T×J^, where *J* is the number of brainstem timeseries. We differentiated for brainstem-to-brainstem and brainstem-to-brain pairs to avoid conditioning on the seed/target timeseries. For a pair of brainstem ROIs *(i,j)*, the covariate matrix included all other brainstem nuclei except the two target regions:

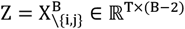

Whereas for brainstem-to-cortex pairs, all brainstem nuclei except the seed region were included:

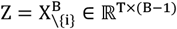

Residuals were obtained from:

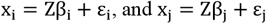

We then estimated partial correlation as:

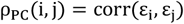

### B. PCA-based Partial Correlation

We implemented PCA-ρ_PC_ within a PC-regression scheme. PCA was applied to obtain the eigen-decomposition of the sample covariance of *Z* matrix (obtained as before) and allowed the reduction of overfitting to noise and collinearity issues by extracting the most informative components. Given the sample covariance matrix of Z (Σ_Z_), first we estimated:

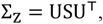

where *U* ∈ ℝ^J×J^ contains the principal axis (loadings) and S is a diagonal matrix of eigenvalues sorted in descending order. We retained the first *nPCs* components, corresponding to the largest singular values (U_nPCs_ = U[: 21: nPCs]). Then the original brainstem covariate matrix was projected onto the reduced orthogonal subspace:

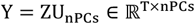

Finally, for each pair *(i,j)*, we regressed out the principal component-derived predictors using MLR and computed PCA – ρ_PC_(*i, j*) by correlating the resulting residuals.

The dimensionality of the PCA conditioning space was determined differently for simulation and experimental data. In the simulation analyses, we used predefined explained variance thresholds in order to assess how PCA dimensionality affects connectome estimation. Conversely, for the experimental fMRI dataset, PCA model order was selected using cross-validation to balance model complexity and generalization performance. We adopted a leave-one-out cross-validation (LOOCV) procedure based on element-wise prediction and minimization of the predicted residual sum of squares (PRESS). Further details are reported in Supplementary Materials.

### C. Regularized Partial Correlation

To compare PCA – ρ_PC_ against commonly used penalized approaches, we implemented L2, L1, and EN partial correlation. Regularization was applied at the regression level to allow a proper comparison with the other implemented solutions. For each pair ∈ {x_i_, x_j_}, we estimated regression coefficients via the penalized optimization problem:

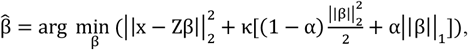

where the first term is the (unregularized) least-square loss, κ ≥ 0 controls the overall penalty strength, and α ∈ (021] sets the mixture between L1 and L2 penalties. In particular, α∼0 corresponds to Ridge, α = 1 to Lasso, and α = 0.5 was used in our implementation for Elastic Net.

After fitting, residual timeseries were computed and used to estimate regularized partial correlation.

Regularization parameters were selected using 5-fold cross-validation and the one-standard-error rule. Further details are reported in Supplementary Materials.

### D. Group-Level Connectomes and Collider effect Correction

We estimated group level connectomes starting from subject-level connectivity matrices. Subject-specific correlation matrices were first Fisher z-transformed, and statistical significance of brainstem–to-brainstem and brainstem–to-brain connections was assessed across subjects using a two-tailed one-sample t-test (*α* = 0.05). To reduce the likelihood of collider-driven false positives, we further combined the partial correlation approaches with pairwise Pearson correlation at the group level, following the rationale proposed in [17], [30]. Specifically, we interpreted the two measures jointly, exploiting the fact that ρ_P_ is ideally unaffected by collider nodes, whereas partial correlation is sensitive to it. Accordingly, a group-level edge was retained in the final corrected connectome only if it was concurrently significant in both ρ_P_ and the corresponding partial correlation-based connectome. Edges significant only in partial correlation but not in ρ_P_ were considered more likely to reflect collider effects and were set to zero. Formally, for each method (**) ∈ {*ρ*_*PC*_, *PCA* − *ρ*_*PC*_, *ρ*_*L*1_, *ρ*_*L*2_, *ρ*_*EN*_}, we defined the corrected group connectome *C*_**_ as:

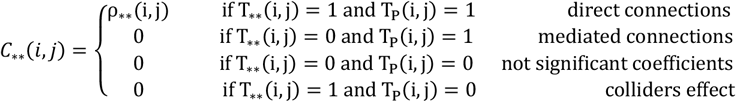

where T refers to the group level binary significance masks (1 = significant, 0 = not significant).

### E. Simulation Study

To benchmark connectivity estimators under controlled conditions, we designed simulations in which the GT network among synthetic timeseries was known and included causal patterns that challenge correlation-based inference, namely confounders, causal chains, and colliders. Simulations were used to progressively evaluate estimator performance across increasingly complex scenarios, ranging from low dimensional toy networks to higher dimensional settings. Details and results on toy examples are reported in Supplementary Materials. For high dimensional settings, we simulated synthetic timeseries using a multivariate linear model [34], [35] as follows:

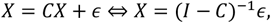

where *ϵ* ∼ *𝒩*(02 *σ*^2^*I*) are independent noise terms. The GT coefficients were defined by a weighted matrix *C* ∈ ℝ^*N*×*N*^ with a zero main diagonal (no self-loops); a non-zero entry *C*_*ij*_ ≠ 0 encodes a direct causal influence *X*_*j*_ → *X*_*i*_, while *C*_*ij*_ = 0 indicates the absence of a direct link between *X*_*i*_ and *X*_*j*_. The presence or absence of a link is defined by the GT networks that we generated as directed acyclic graphs (DAGs) under the Erdős–Rényi model (ER-DAGs). The network structure was set up by drawing a random permutation of the N nodes and allowing an edge i→j only when node i precedes node j in the permutation. Because of this “forward-only” constraint, the resulting entire graph was acyclic by construction. Given a target edge count *M*, we then sampled exactly *M* edges uniformly from the set of admissible ordered pairs. We varied the total number of nodes *N* ∈ [202 352 502 802 1202 1802 2302 3002 3452 3802 5002 600], and defined the number of brainstem-like nodes as *N*_*BR*_ ∈ [4, 8, 11, 18, 26, 40, 51, 66, 76, 84, 110, 132], respectively. The number of non-zero edges (*M*) was controlled by fixing the network density (*D=5%*) for each *N* and corresponding *N*_*BR*_. Specifically, after defining the first *N*_*BR*_ nodes as brainstem-like, M was set separately for the brainstem-to-brainstem subnetwork and for the remaining nodes as 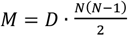. To approximate the temporal properties of rs-fMRI, simulated signals were band-pass filtered ([0.01,0.1] Hz).

For each realization of *ϵ*, a single-subject dataset X was simulated. We then generated 25 synthetic datasets to emulate multiple subjects by resampling the noise term while keeping the network structure C fixed and varying the weight of the existing connections. Each edge weight was randomly sampled from a Gaussian distribution centred on its corresponding GT value in C, with a standard deviation equal to 0.4. Thus, between-subject variability reflected not only stochastic noise but also biological variability. We designed simulations to evaluate how GT recovery depends on key parameters. In addition to *N* and *D*, we varied the number of timepoints *T*, the noise level *σ* (to modulate signal-to-noise ratio), and the explained variance threshold V used to select the number of components in *PCA* − *ρ*_*PC*_. To assess the effect of each parameter, we varied one parameter at a time while keeping the others fixed. The reference configuration was T = 800, *σ* = 0*N*01, N = 300, V = 80%, D = 5%. The tested ranges were as follow: T ∈ [4002 6002 8002 100021200], *σ* ∈ [0.01, 0.05, 0.1, 0.5, 1], and *V* ∈ [0.7, 0.8, 0.9, 0.95, 0.99].

For each of the 25 repetitions, we estimated *ρ*_*P*_, *ρ*_*PC*_, and *PCA* − *ρ*_*PC*_ with and without collider correction. Of note, to replicate the conditioning strategy adopted for the empirical data, we selected a subset of nodes in the simulated networks, which we defined as “brainstem-like” and used as conditioning variables in the partial correlation framework. We quantified edge-level metrics including FNR, FPR, TPR, and BA (defined as the arithmetic mean between sensitivity and specificity).

We evaluated these metrics on the connections from the brainstem-like nodes. In Supplementary Materials, we report additional simulation analysis in which we aggregated the connectivity matrices estimated across the 25 repetitions and replicated the group level analysis.

### F. Experimental fMRI Dataset and Processing

We analysed a publicly available resting-state fMRI dataset from the OpenNeuro repository (accession number ds003673; DOI: 10.18112/openneuro.ds003673.v2.0.1), including 27 healthy adults scanned on a Siemens 3T MAGNETOM Prisma system at the Yale Magnetic Resonance Research Center. Each subject underwent one T1-weighted anatomical acquisition and two resting-state fMRI runs acquired within the same session. Structural images were acquired using a 3D T1-weighted MPRAGE sequence (TR = 2400 ms, TE = 1.22 ms, flip angle = 8°, 1 mm isotropic resolution), whereas resting-state fMRI data were acquired using a multiband gradient-echo EPI sequence (TR = 1000 ms, TE = 30 ms, flip angle = 55°, multiband factor = 5, voxel size = 2 × 2 × 2mm, 75 slices, 410 volumes per run). For further details on the data and acquisitions refer to [36].

Functional and anatomical preprocessing were performed in AFNI. Anatomical images were skull-stripped and nonlinearly normalized to MNI space. Functional preprocessing included despiking, slice-timing correction, rigid-body motion correction, co-registration to the anatomical image, spatial normalization, spatial smoothing using a 4 mm FWHM Gaussian kernel, voxel-wise scaling to percent BOLD signal change, nuisance regression, and temporal band-pass filtering (0.01–0.12 Hz). All spatial transformations were concatenated and applied in a single interpolation step to minimize resampling-related artifacts. Nuisance regression was performed within a generalized linear model framework and included: (i) six rigid-body motion parameters and their temporal derivatives, (ii) average white matter and cerebrospinal fluid signals extracted from eroded nuisance masks, and (iii) third-order polynomial trends to account for low-frequency signal drift. Temporal filtering was performed simultaneously with nuisance regression in order to avoid frequency reintroduction artifacts. Runs failing motion- or signal-based quality control (QC) criteria were excluded (see Supplementary materials for details). After QC, 16 participants (two runs each) were retained for further analysis.

Brainstem ROIs were defined using the Brainstem Navigator probabilistic atlas, including bilateral and midline nuclei in MNI space. Cortical targets were defined using the Glasser multimodal atlas projected to volumetric space. For each ROI, representative BOLD timeseries were obtained by averaging gray matter voxel signals within the corresponding parcel, ROIs timeseries from the two runs were concatenated prior to FC estimation with *ρ*_*P*_, *ρ*_*PC*_, and *PCA* − *ρ*_*PC*_ and regularized *ρ*_*PC*_. For all the methods, we focused on brainstem-to-brainstem and brainstem-to-brain FC (i.e., on the submatrix ρ ∈ ℝ^B x (B+CTX)^).

### G. Multicollinearity and Stability assessment

Multicollinearity occurs when the predictors in a regression model are strongly correlated so that they carry redundant information about the outcome variable and make the estimation of individual regression effects unstable [20], [37]. In this setting, the ordinary least square solution becomes numerically ill-conditioned because it depends on (*X*^*T*^*X*) ^−1^, which may be nearly singular when columns of X close to linearly dependent (i.e., geometrically lie on the same hyperplane). An ideal regression design involves orthogonal predictors, which maximizes the precision of coefficient estimates and optimizes the model’s power [38]. Since the PCA, while reducing dimensionality, maps the set of regressors into an orthogonal (uncorrelated) component space, it provides a direct and effective strategy to mitigate multicollinearity issues [39]. Regularization approaches can also improve numerical stability by shrinking coefficients, especially under strong predictor correlations [40], [41], [42].

To quantify multicollinearity, for each subject, we computed the condition numbers (*n*) of the standardized regressors covariance matrix *C*_*z*_ as:

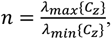

where *λ*_max_ and *λ*_min_ are the largest and smallest eigenvalues, respectively.

Specifically, for naïve *ρ*_*PC*_, *n* was evaluated in the original regressor space; for *PCA* − *ρ*_*PC*_ the condition number was computed in the PCA-transformed regressor space after dimensionality reduction (i.e., using the retained principal component scores as regressors); for *ρ*_*L*2_, we computed a penalized conditioning metric based on the regularized matrix (i.e., *n*{*X*^*T*^*X* + *κI*}); whereas for *ρ*_*L*1_ and *ρ*_*EN*_, which are methods inducing sparsity, we calculated *n* on the active set (i.e., the submatrix of regressors with non-zero estimated coefficients).

For each method, we then summarised these values across subjects using median, interquartile range (IQR), and median absolute deviation (MAD).

Additional analyses on the assessment of multicollinearity and stability are reported in Supplementary Materials.

## Results

### A. Synthetic Data

We evaluated the behaviour of the different FC estimators in large scale simulations designed to mimic realistic connectomic settings. Figure 1 shows FPR, FNR, TPR, and BA as a function of the number of regions, reported as median values across simulations together with their interquartile range (IQR). A figure reporting the effect of varying the number of timepoints is reported in Supplementary Materials. No substantial effects were observed when varying the amount of explained variance retained in PCA and the signal-to-noise ratio of the timeseries (data not shown).

**Figure 1.**
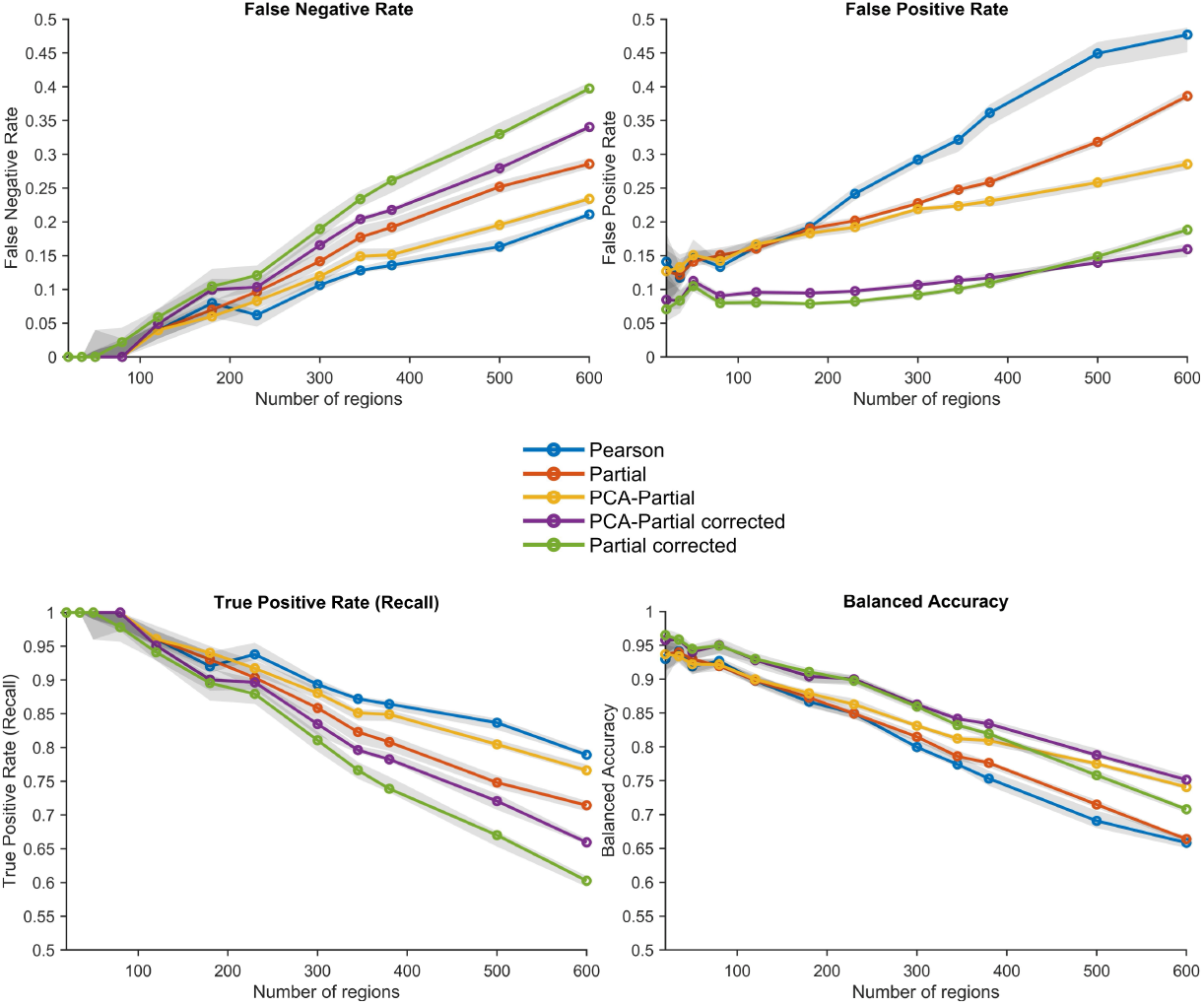
Effect of network dimensionality. Performance of FC estimators used in this work as a function of the number of regions included in the network. The panels report false negative rate (FNR), false positive rate (FPR), true positive rate (TPR/recall), and balanced accuracy (BA). Solid lines indicate median values across simulations, and shaded areas represent interquartile ranges.

As the number of regions increased, the recovery of the GT network became progressively more challenging. This effect was accompanied by a corresponding increase in collinearity within the conditioning set. Across simulations, ρ_P_, ρ_PC_, and PCA − ρ_PC_ showed comparable performance up to approximately 200 regions (with differences shown between these and the collider-corrected versions). Beyond this point, estimators’ performance diverges. ρ_P_ consistently showed high sensitivity (TPR), but this was associated with a rapidly increasing FPR, substantially higher compared to other methods, due to the accumulation of numerous indirect or mediated connections. ρ_PC_ substantially reduced false positives by conditioning on the brainstem-like nodes, resulting in increased specificity. However, FPR still increased with network dimensionality, likely due to the growing collinearity among regressors. This gain in specificity was accompanied by an increase of the FNR, particularly in larger networks. PCA-based partial correlation achieved a more balanced trade-off between sensitivity and specificity. By reducing collinearity in the conditioning set, it limited the increase in false positives while avoiding the strong loss of sensitivity observed in standard partial correlation, resulting in improved balanced accuracy compared with both ρ? and ρ_PC_. Finally, both ρ_PC_ and PCA – ρ_PC_ showed consistent improvements when combined with collider correction. While this correction led to a slight increase in FNR and a corresponding decrease in TPR, it produced a clear reduction in FPR. This pattern roughly follows that observed for the uncorrected versions, ultimately resulting in slightly better performance of the collider-corrected PCA – ρ_PC_ method across all methods.

### B. 3T rs-fMRI Data

For each subject, we computed the condition number (*n*) of the standardized regressor covariance matrix and summarised the results pooled across runs using median, interquartile range (IQR), and median absolute deviation (MAD). As shown in Table 1, standard *ρ*_*PC*_ was associated with marked ill-conditioning, consistent with strong multicollinearity among brainstem covariates. Regularized methods mitigated ill-conditioning by reducing *n* compared to *ρ*_*PC*_; however, condition numbers remained substantially higher than those observed in the PCA-transformed space. This indicates that PCA-based decorrelation and dimensionality reduction effectively stabilised the estimation problem. We finally evaluated whether the proposed framework yielded physiologically plausible connectivity patterns. Figure 2A reports group level connectomes for the superior colliculus (SC) estimated using *ρ*_*P*_ and collider-corrected *PCA* − *ρ*_*PC*_. Left sides display statistically significant cortical connections projected onto cortical surfaces for visualization purposes. Right sides shows circular plots of significant connections with brainstem nuclei. Both brainstem-to-brain and brainstem-to-brainstem results are presented after averaging left and right homologous regions for both seeds and targets, resulting in symmetric connectomes. PCA-based partial correlation reduces the contribution of mediated or indirect interactions from other brainstem nuclei, yielding markedly sparser and anatomically structured connectomes compared to the corresponding Pearson maps. At the cortical level, fewer but more focal connections emerge, with connectivity patterns appearing more topologically distributed and spatially organized. Clusters of connections focus on functionally relevant regions according to the role of each nucleus and highlight a clear differentiation between nuclei-specific patterns. Within brainstem FC, the differences between methods are even clearer. The circular plot obtained using *ρ*_*P*_ shows unspecific connectivity patterns that are widespread across brainstem nuclei and include connections involving areas from various functional systems. In contrast, *PCA* − *ρ*_*PC*_, while mitigating spurious correlations, identifies more selective connectivity patterns, with connections primarily targeting nuclei belonging to the same functional pathway as the seed region. Specifically, for SC cortical pathways focused on occipital regions, including connectivity with early and intermediate visual cortices. Differences between methods are more evident at the brainstem level. The circular plots show that *ρ*_*P*_ produces a more globally dense connectivity pattern, whereas *PCA* − *ρ*_*PC*_ emphasizes interactions with orienting-related and reticular nuclei (e.g., PAG, mRT, ISRT). Interestingly, as highlighted in Figure 2B, the subcortical interactions of SC are characterized by short-range anatomical distances.

**TABLE I.** Conditioning of the brainstem Covariate Matrix across Methods.

|  | *Partial*<br>*Correlation* | *PCA*<br>*Regularization* | *L1 Penalty* | *L2 Penalty* | *EN Penalty* |
| --- | --- | --- | --- | --- | --- |
| Median<br>± IQR | 524.24 ± 318.71 | 7.72 ± 12.06 | 174.95 ± 129.33 | 177.31 ± 143.14 | 181.83 ± 134.32 |
| MAD | 217.23 | 12.07 | 11.08 | 106.23 | 11.48 |

Condition numbers of the standardized brainstem regressor covariance matrix are summarized across subjects using the median with interquartile range (IQR) and median absolute deviation (MAD). Values refer to data concatenated across run 1 and run 2. $PCA - \rho_{PC}$ was computed in the PCA-transformed space; Ridge was computed via the L2-penalised regression formulation; Lasso and Elastic Net were computed on the active set.

**Figure 2.**
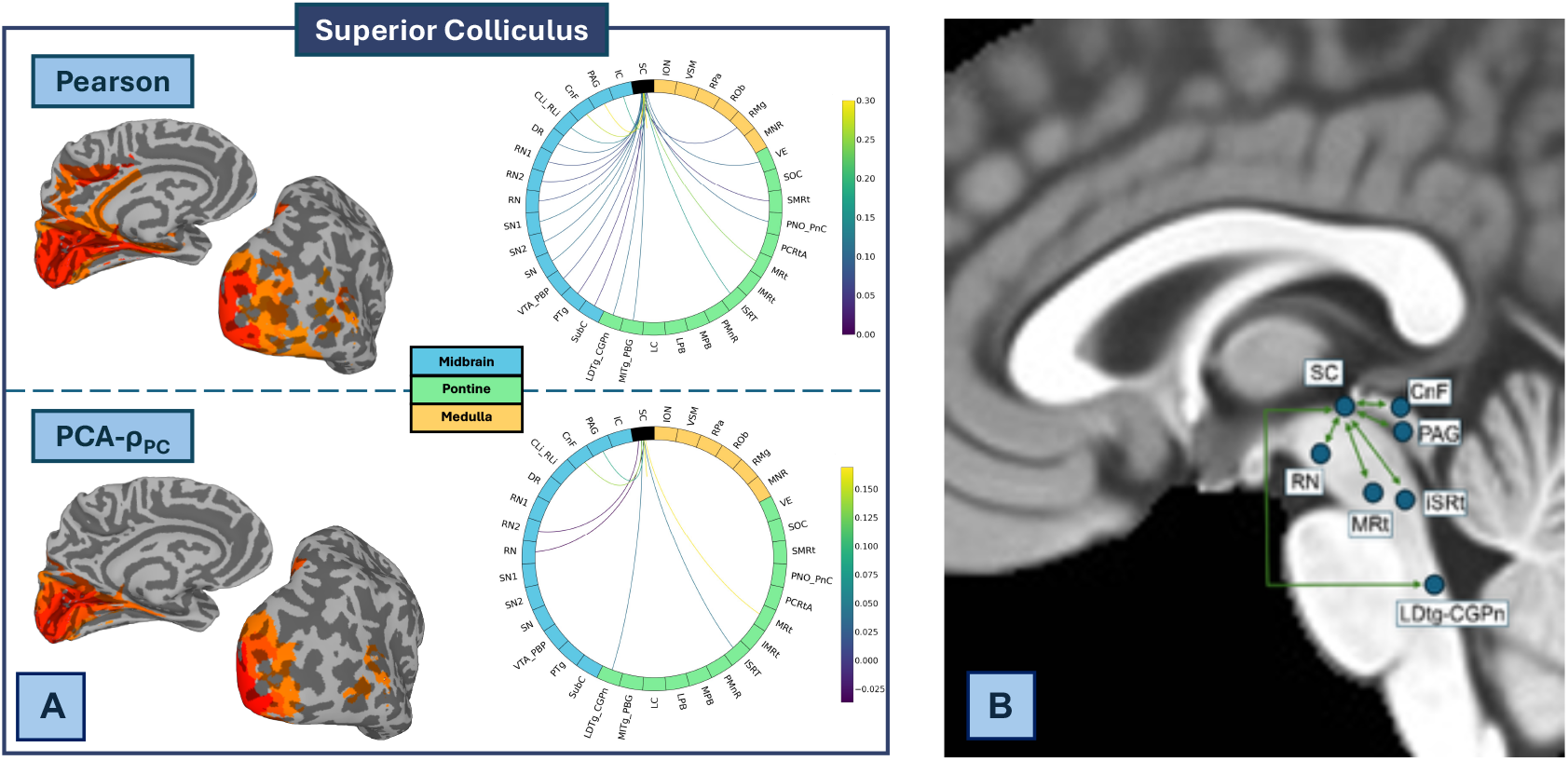
Functional connectomes of the Superior Colliculus. A) Cortical and subcortical connectomes obtained with Pearson and PCA-based partial correlation. On the left, the statistically significant cortical connections projected onto the cortical surface are reported; on the right circular plots of significant connections between each nucleus and other brainstem nuclei are shown. In PCA-based partial correlation the regions with statistically significant correlations with SC are the following: primary visual cortex, medial superior temporal area, second visual area, third visual area, area 45, are 47 lateral, area anterior 47r, presubiculum, prostriate area, dorsal transitional visual area, area FST, ventromedial visual area 2. B) Circuit of the subcortical interactions (PCA-based Partial Correlation).

The connectome of the inferior colliculus (IC) is reported in supplementary materials, revealing that *PCA* − *ρ*_*PC*_ isolates a sensorimotor and multisensory functional organization, different from the broad multimodal covariance structure captured by *ρ*_*P*_.

## Discussion

In this work, we proposed and evaluated a *PCA* − *ρ*_*PC*_ framework specifically tailored to the estimation of brainstem direct FC from rs-fMRI. In both synthetic and experimental data, our approach consistently yielded sparser and more structured connectomes compared with *ρ*_*P*_, while mitigating the instability associated with naïve *ρ*_*PC*_ under ill-conditioned regression settings. Overall, our results suggest that PCA-based conditioning provides a useful compromise between the high sensitivity but limited specificity of *ρ*_*P*_ and the unstable estimation regime of standard *ρ*_*PC*_. This framework was motivated by two major methodological and neurophysiological challenges. First, as also recently observed in [15], brainstem nuclei timeseries share strong multicollinearity, which is quantified in our study with the high condition numbers of the regression design matrices. Shared anatomical inputs, common neuromodulatory influences, and the intrinsically dense topology of brainstem systems likely contribute to this behavior. Under these settings, conditioning and reliability analyses prove that standard *ρ*_*PC*_ becomes highly sensitive to noise and inversion instability. By introducing PCA regularization, our framework stabilized the regression problem while preserving the dominant covariance structure of the brainstem network. A second key issue concerns the interpretation of functional associations in highly interconnected systems. The comparison between *ρ*_*P*_ and partial-correlation-based approaches, in both simulations and experimental data, highlighted false positive interactions driven by confounders, indirect pathways, and shared network fluctuations, which are substantially attenuated after conditioning on the brainstem covariate. We did not implement a fully unconstrained multivariate estimation across all brain regions; instead, we adopted a physiologically motivated choice and restricted the conditioning space to the brainstem nuclei themselves. The real data analyses prove that this choice preserves the interpretability of the resulting estimates. Under this formulation, the retained principal components can be interpreted as capturing dominant patterns of shared activity within the brainstem network, while the residual associations quantify the contribution of each nucleus after accounting for the variance shared with the remaining brainstem nuclei. The resulting connectivity estimates retain a nucleus-centered interpretation and are more directly related to the specific functional role of individual nuclei within large-scale brain interactions.

### A. Synthetic Data

Simulations results demonstrate that the previously discussed differences arise from the intrinsic properties of the estimators. We replicated the same conditioning strategy in simulations where only a set of the synthetic nodes were considered as brainstem-like. Simulations further clarified the conditions under which *PCA* − *ρ*_*PC*_ becomes more advantageous. Under relatively low-dimensional (up to 200 regions) and weakly collinear settings, *ρ*_*P*_, *ρ*_*PC*_ and *PCA* − *ρ*_*PC*_ showed overall comparable performances; differences are shown between these and collider-corrected versions. In high network dimensionality, Pearson correlation maintained high sensitivity but captured more indirect interactions leading to inflated FPR. Standard partial correlation reduced FP, indicating improved specificity when conditioning on brainstem-like nodes, but this gain was accompanied by reduced sensitivity and only limited improvement in balanced accuracy. By contrast, *PCA* − *ρ*_*PC*_, by decorrelating the conditioning set, achieved a more favourable trade-off between sensitivity and specificity: while maintaining FNR and TPR approximately comparable to those of *ρ*_*P*_, it markedly improved FPR, which translate into higher BA in larger networks. Summary analyses in supplementary materials confirmed this pattern, above all in the most challenging settings, namely when network density was high, collinearity was strong, or the dimensionality of the system increased. Importantly, these conditions are not uncommon in resting-state fMRI connectomics. Modern cortical and subcortical atlases frequently include several hundred regions, while the number of temporal samples remains inherently constrained by acquisition duration and temporal resolution. In addition, preprocessing operations such as temporal filtering and spatial smoothing, together with shared physiological and neuromodulatory fluctuations, can substantially increase correlations among regional timeseries, leading to ill-conditioned regression problems even in moderately low dimension networks. More generally, similar problems occur in several multivariate biomedical applications, including multimodal neuroimaging integration, and high-dimensional physiological or clinical modeling.

For both *ρ*_*PC*_ and *PCA* − *ρ*_*PC*_, collider correction further reduced FP, although FNR increased and TPR decreased. This indicated that correction of collider-induced associations may lead to an overcorrection in tightly coupled network structures where several motifs interact simultaneously. Moreover, when multiple causal motifs coexist and nodes participate simultaneously in the information exchange within the network, closed loops and densely interconnected subnetworks occur. In such cases, the shared variance across regions makes different nodes of the network have synchronized timeseries, and, if only a subset of relevant variables is included in the conditioning process, the effects of partialization become less intuitive. Toy examples reported in supplementary materials further exemplified such scenarios. When selecting variables for correlation-based analyses, either in connectomic studies or in broader multivariate scenarios, researchers often implicitly assume simplified interaction models among variables, such as the canonical causal motifs illustrated in Supplementary Figure 1. However, real world networks rarely consist of isolated motifs. Multiple interaction patterns can coexist and interact within the same network, and the set of variables included in the conditioning set often represents only a subset of the potential sources of variability influencing the system.

### B. 3T fMRI Data

Simulation results represent a scaffold to interpret connectomes from real fMRI data. Here, our findings demonstrate that *PCA* − *ρ*_*PC*_ provides a robust and more interpretable estimate of FC in a challenging setting, namely the brainstem networks characterized by high interdependence between nuclei, low SNR, and strong shared physiological variance. *PCA* − *ρ*_*PC*_ allowed us to isolate brainstem direct functional relationships while maintaining high stability. In line with a recent methodological work highlighting the benefits of regularization for improving FC robustness and reliability [26], our findings suggest that PCA-based regularization can stabilize multivariate estimates not only by reducing overfitting to noise, but also by improving the conditioning of the regression problem. In fact, a first major finding is that *PCA* − *ρ*_*PC*_ directly addresses the multicollinearity problem that limits the application of naïve *ρ*_*PC*_. The condition numbers associated with the unregularized *ρ*_*PC*_ were systematically high, indicating that the regression problem underlying residualization was often severely ill-conditioned. This implies that even small perturbations in the data can produce large changes in regression coefficients that reflect in edge connectivity weights. This interpretation is supported by the run-to-run supplementary analyses, where standard *ρ*_*PC*_ showed reduced pattern reliability, larger run-to-run edgewise change, and a subset of subjects with near-saturated correlations. *PCA* − *ρ*_*PC*_ directly targets the source of this, resulting in orders of magnitude improvements in conditioning and in more stable connectome estimates, while still preserving the multivariate logic of partial correlation. The comparison with other regularized estimators helps position *PCA* − *ρ*_*PC*_ within the broader literature on direct FC estimation [23], [25], [43], [44], [45]. In line with previous works, L1, L2, and EN regularized partial correlation all improved some aspects of stability relative to standard partial correlation. In particular, *L*2 − *ρ*_*PC*_ yielded the highest run-to-run reproducibility, closely matching *ρ*_*P*_. However, this advantage must be interpreted carefully. Ridge stabilizes the regression coefficients through shrinkage, but it does not explicitly decorrelate the design matrix, therefore alone it does not fully resolve the underlying conditioning problem. As a possible consequence, the resulting connectomes are comparable with *ρ*_*P*_ maps, suggesting that greater run stability may come at the cost of reduced specificity in isolating conditional dependencies. Sparse penalties, in contrast, did not perform as well in this dataset in terms of run-to-run reproducibility, likely because instability in the active set can amplify run- and subject-level variability when predictors are strongly collinear. Taken together, these findings suggest that *PCA* − *ρ*_*PC*_ occupies a useful intermediate regime: less stable than simple correlation, but substantially more robust than unregularized partial correlation, while retaining a clearer multivariate interpretation than shrinkage-based solutions whose connectomes remain closer to full correlation.

Beyond numerical stability, the real data analyses demonstrate that *PCA* − *ρ*_*PC*_ does not simply reduce connectivity density but reveals a qualitatively different organization of brainstem functional networks. Under *ρ*_*P*_, the nuclei appear embedded in a globally interconnected system, with widespread cortical and subcortical coupling. In contrast, *PCA* − *ρ*_*PC*_ preserves a more constrained subset of interactions and reveals distinct connectivity profiles, supporting a model in which the brainstem is organized into partially segregated functional circuits [16], [46]. This effect was particularly evident at the brainstem level, where *ρ*_*P*_ identified extensive coupling across nuclei belonging to sensory, motor, autonomic, and arousal-related systems. Such dense networks are broadly compatible with the known high interconnectivity of the brainstem, but the inability to distinguish direct from mediated couplings limits a straightforward physiological interpretation.

### C. Neurobiological Interpretability

We interpreted our real data connectomes in the light of known neuroanatomical brainstem organization. Under *ρ*_*P*_, SC and IC showed strong and diffuse couplings, while the selectiveness of *PCA* − *ρ*_*PC*_ revealed nucleus-specific connectivity that might indicate potential fingerprints. Importantly, these last connectivity patterns are more consistent with known neuroanatomical organization. The SC exemplified this effect particularly clearly. *ρ*_*P*_ map includes extensive coupling with higher-order visual, parietal, and limbic regions. This broad multimodal connectivity pattern may reflect large-scale network co-fluctuations and likely overestimates the extent of direct SC interactions. After conditioning, *PCA* − *ρ*_*PC*_ isolates a more constrained plausible network focused on early visual processing, motion sensitivity, and sensorimotor integration. Specifically, it showed statistically significant correlations with early and intermediate visual cortices (V1–V3), motion-sensitive regions (e.g., FST), and peripheral vision–related cortex as oculomotor areas, extending to spatial representations in the presubiculum and to frontal regions (areas 45/47). At the subcortical level, SC was coupled with midbrain nuclei involved in orienting and sensorimotor coordination, including PAG, cuneiform nucleus (CnF), reticular formations (MRt, ISRT), LDTg, and RN. These results highlight a visuomotor network in which the SC acts as a hub integrating visual inputs with motor and spatial representations to guide rapid, action-oriented behaviours. This is in line with known circuits supporting orienting responses, locomotion, and behavioural state regulation [47], [48], [49]. Moreover, the selective FC with early visual and motion-related regions supports the view of SC as a hub of sensory-driven and action-oriented processes, rather than a node widely related to cognitive systems [50], [51].

### C. Implications for brainstem functional connectivity studies

The brainstem remains relatively understudied in fMRI due to technical and methodological challenges, particularly in the context of FC and large-scale functional organization. The discussed methodology and findings have broader implications for brainstem FC studies and can advance further analyses, especially correlation-based ones, that aim to investigate investigating neuromodulatory systems, sensory integration, and brainstem contributions to large scale brain networks in both healthy and pathological conditions.

Importantly, because conditioning was restricted to brainstem nuclei, the resulting connectomes are not intended to represent fully “direct” whole-brain networks in the strict graphical model sense. Indirect effects mediated through cortical or subcortical regions may still be present. This design choice reflects the trade-off of focusing on the contribution of specific brainstem nuclei over the elimination of all possible mediated pathways and also align with the main neuroscientific question of the study “how do individual brainstem nuclei contribute to interactions within the brainstem and with the other whole-brain regions?”. On the other hand, the brainstem-only conditioning strategy also suggests a natural methodological extension. Future work could adopt hierarchical or iterative conditioning schemes, in which subsets of subcortical and cortical parcels are progressively introduced into the conditioning set, thereby reconstructing complete ascending functional pathways.

### D. Methodological considerations and limitations

The reduction in FC density observed with *PCA* − *ρ*_*PC*_ is consistent with the attenuation of indirect and mediated interactions that, in the simulations, contributed substantially to the FPR of *ρ*_*P*_. At the same time, simulations also showed a partial increase of FNR, which may be reflected in missing direct functional connections in the real data *PCA* − *ρ*_*PC*_ connectomes. Taking into account the surplus of significant *ρ*_*P*_ connections over the *PCA* − *ρ*_*PC*_’s ones together with the rational underlined the collider correction arise the possibility that Pearson PCA-based partial correlation approaches may complement each other in an effort to unravel the pattern of direct vs indirect connections among brain areas. By systematically varying the conditioning set (e.g., exploiting a recently developed algorithm to perform the conditioning strategy [23]) it may also become possible to explore how specific subsets of regions contribute to the observed interactions, offering insights into the identification of potential confounders, collider structures, or causal chain elements. Importantly, this perspective is not intended to replace formal causal modelling approaches but, as mentioned in a recent work [30], it may provide a first step and data-driven scaffold for generating targeted hypotheses about the organization of complex brain networks to be tested with directional or causal modelling approaches, such as Dynamic Causal Modelling or Structural Equation Modelling.

The improved specificity of *PCA* − *ρ*_*PC*_ and lower edge density should not be interpreted only as a methodological consequence. More selective and anatomically constrained connectomes may be advantageous not only for network interpretation [52], [53], but also for extracting more specific and constrained features for translational applications, including classification, prediction, and machine-learning or deep-learning frameworks.

Some limitations should be considered. First, the analyses relied on an instantaneous, zero-lag FC model, which is a common choice in rs-fMRI, particularly given the relatively low temporal resolution, but it does not explicitly model delayed interactions. Extending the framework to lagged measures would be an important next step, especially in light of the hemodynamic delays inherent in fMRI and the response profiles of brainstem and subcortical nuclei which differ from cortical areas themselves. Second, we used 3T data that may have limited sensitivity for the smallest nuclei. In fact, although 2 mm isotropic resolution is common in 3T fMRI studies, some nuclei are characterized by cross-sectional dimensions comparable to or smaller than a single voxel (e.g., locus coeruleus and raphe nuclei). The measured BOLD signal may, therefore, be affected by partial-volume. For these reasons, we focused the physiological interpretation on the SC and IC, whose dimensions are relatively compatible with the spatial resolution of this dataset. Furthermore, a 4 mm FWHM Gaussian smoothing kernel was selected as a compromise between improving SNR and limiting excessive spatial blurring across adjacent nuclei. Several brainstem studies have used 7T acquisitions [15], [54], [55], [56]; however, some analyses which demonstrate the high translatability of brainstem FC results from 7T to 3T scanners have already been performed [57], [58]. We focused on 3T data, which remain the standard in most research and clinical settings and strengthen the translational relevance of the approach.

### E. Conclusions

Our findings support *PCA* − *ρ*_*PC*_ as a useful tool to investigate brainstem direct FC. By combining a brainstem-focused conditioning strategy with PCA decorrelation and collider correction, our framework reduces multicollinearity, improves robustness, and yields nucleus-specific connectivity patterns that are more readily interpretable. Specifically, *PCA* − *ρ*_*PC*_ can be viewed as a bridge between dense *ρ*_*P*_ maps and selective direct FC, offering a practical framework for studying highly collinear networks in rs-fMRI. More broadly, the proposed framework could be extended to task-based designs, where distinguishing shared state-dependent variance from more specific circuit-level interactions may be equally important. Such an approach may be especially valuable in clinical conditions involving brainstem dysfunction, as well as in neuroscientific contexts focused on cognition, prediction, memory, or arousal-related processing. Finally, the potential utility of this strategy may extend beyond fMRI FC alone. Similar principles may find application in a wide range of multivariate neuroimaging applications, spanning from other modalities to multimodal integration frameworks (e.g., fMRI, EEG, pupillometry, or other complementary signals).

