## supplementary materials for "Physiologically Informed PCA-Partial Correlation for highly Collinear Brainstem fMRI Networks"

### 1. SCHEMATIZATION OF CANONICAL CONNECTIVITY PATTERNS

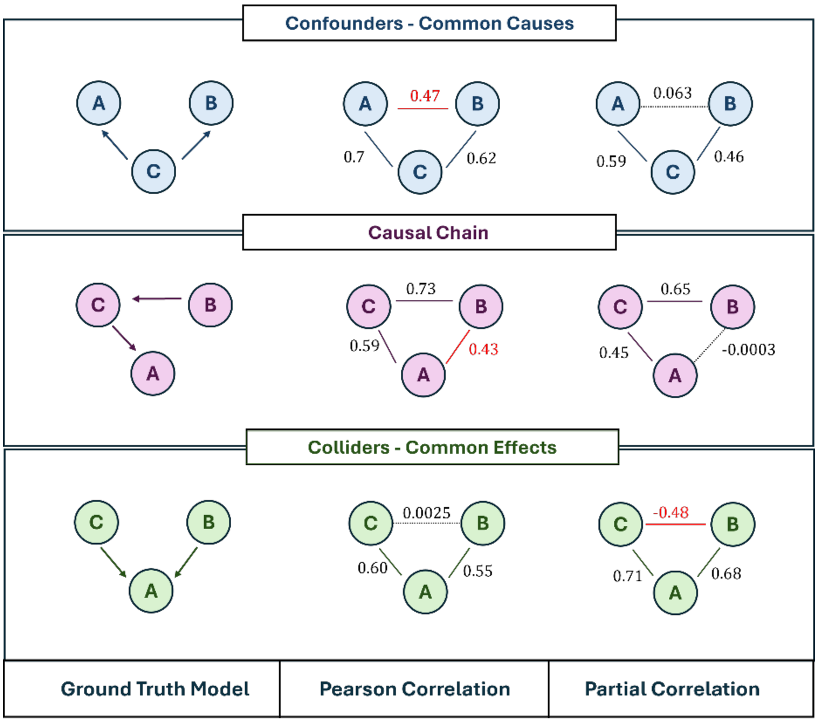


*Supplementary Figure 1: Canonical connectivity patterns that bias functional connectivity estimates. Common examples are shown as ground truth model (left), Pearson correlation (middle), and partial correlation (right). A) Confounder/common cause with Pearson leading a non-zero interaction despite the absence of a direct link. B) Causal chain with Pearson resulting in A-B association mediated by C. C) Collider/common effect for which partial correlation, by conditioning on C, can induce spurious dependency. Timeseries were estimated with the model employed for simulation.*

### 2. SELECTION OF MODEL PARAMETERS

Model parameters were selected differently depending on the regularization approach. Pearson correlation and naïve partial correlation do not require hyperparameter tuning. In contrast, PCA-, L2-, L1-, and EN- $\rho_{PC}$ require parameter selection for optimal performance. To this aim, we adopted cross-validation and implemented two robust and validated strategies tailored to the specific regularization scheme.

For PCA-Partial correlation, the relevant tuning parameter is the dimensionality of the PCA conditioning space. For the experimental fMRI dataset, PCA model order was selected using cross-validation to balance model complexity and generalization performance. We adopted a leave-one-out cross-validation (LOOCV) procedure based on element-wise prediction, as described in [1]. Specifically, for each subject, we applied LOOCV to the mean-centred brainstem ROI time series. At each LOOCV fold, one timepoint was held out as a test observation, and PCA was fitted on the remaining (T-1) samples. For a candidate model order $k$, we assessed how well the PCA model could reconstruct the held-out observation using a leave-one-variable-out procedure: each variable $j$ in the test vector was predicted in turn while treating that variable as missing [1]. Concretely, test scores were estimated by least squares using the remaining (J-1) observed variables and the corresponding subset of the training loading matrix (computed via Moore–Penrose pseudoinverse). The left-out variable was then reconstructed from the estimated scores and the loading matrix. For each pair $(t, j)$, the prediction error was computed as the squared difference between the observed and reconstructed values. Errors were summed across all variables and LOOCV folds to yield the predicted residual sum of squares (PRESS) as a function of $k$. The optimal dimensionality (*nPCs*) was chosen as the value of $k$ that minimized the cross-validated PRESS.

Regularized partial correlation approaches require the tuning of the penalty parameter κ, which determines the strength of regularization: larger values impose stronger constraints, leading to sparser solution for L1 or increased shrinkage for L2, at the cost of increased prediction error (MSE). We selected κ using K-fold cross-validation (K=5) [2] as a compromise between bias and variance [3], [4]. Within each fold, we evaluated a geometric sequence of N=100 candidate κ values, where the largest value corresponded to the smallest penalty for which the null model is obtained (all regression coefficients equal to zero, $\hat{\beta} = 0$ ). For each candidate κ, the model was fitted on K-1 partitions and evaluated on the held-out fold using penalized MSE, producing a cross-validated error curve as a function of κ. In order to favour stable solutions and reduce sensitivity to noise [2], we selected $\kappa_{1SE}$ according to the "one-standard-error" rule: among candidate values, we chose the largest κ whose cross-validated error was within one standard error of the minimum.

### 3. SIMULATION STUDY

Simulations were used to progressively evaluate estimator performance across increasingly complex scenarios, ranging from low dimensional toy networks to higher dimensional settings. Specifically, we aimed to investigate: (i) the qualitative behaviour of correlation estimators in the presence of interacting causal motifs; (ii) the accuracy of GT network recovery in high dimensional networks; and (iii) the effect of restricting the conditioning set to a subset of regions while leaving the remaining nodes latent.

#### 3.1. Toy example Networks

In this section, we report the analysis performed on the Toy Example networks, the effect of varying the number of timepoints in high dimensional simulations, and the summary analyses across repetitions of simulations.

For the toy examples, we first generated GT networks by manually specifying the GT connectivity structure (i.e., the presence or absence of edges) to enforce the coexistence and interaction of multiple causal motifs within the same network. Given the network, coefficient values were assigned by sampling non-zero weights from a uniform distribution on [-1,1]. To avoid near-zero effects, coefficients in [-0.1,0) were truncated to -0.1 and those in (0,0.1] were set to +0.1. We then generated timeseries following the same model used in the high dimensional settings. We built 5 networks, each including 11 nodes. The first network, in addition to simple one-to-one connections, contained two causal chains; the second contained one collider and one confounder. We then simulated two additional examples that included a subnetwork embedded within a cyclic structure: one composed of a collider, a confounder and a causal chain, and another including three consecutive causal chains. Finally, the fifth network consisted of two non-interacting subnetworks. Three of the five networks included four nodes included in the conditioning set (i.e., brainstem-like nodes), whereas the remaining two included five. For each example, we estimated $\rho_P$, $\rho_{PC}$, $PCA - \rho_{PC}$, and subsequently applied the collider correction. The GT structures and the estimated connectivity matrices are reported in figure 2 (note that $\mathrm{PCA} - \rho_{PC}$ results are not shown here because, in these low dimensional settings without multicollinearity, they gave results identical to those obtained with $\rho_{PC}$). For each network, the GT structure is shown in the upper panel, whereas the matrices estimated using $\rho_P$, $\rho_{PC}$ and its collider-corrected version are reported in the lower panel.

Across the toy examples, the qualitative behaviour of the estimators when multiple interaction patterns coexist within the same network remains consistent with the well-known simplified scenarios illustrated in Fig.1. Pearson correlation, due to its bivariate nature, recovers dense networks and systematically identifies a large number of connections when the GT is characterized by causal chains and confounders. These mediated associations appear as additional edges in the estimated connectomes. Examples of such false positives (FP) can be observed between non-adjacent nodes in fig. 2A and fig. 2E, as well as between nodes 1–4 in fig. 2B. In contrast, $\rho_{PC}$ largely removes these indirect associations after partialization. This is evident, for example, in the subnetwork involving nodes 8-11 in fig. 2E. Two additional findings emerge from the $\rho_{PC}$ results. First, in our simulations, only the green nodes (considered as brainstem-like nodes to match the real fMRI data framework) were regressed out. As a result, some FP (indirect) connections are still found in the $\rho_{PC}$ estimates (e.g., between nodes 2-3 in fig. 2A or between nodes 1-4 in fig. 2E), reflecting the composition of the conditioning set. Second, $\rho_{PC}$ remains sensitive to collider-induced associations and reveals FP that are not present in $\rho_P$ estimates. Examples of these interactions are evident in fig. 2E,where the FP between nodes 2-3 is driven by the collider effect caused by node 1, and in fig.2B. These examples show situations in which conditioning can introduce spurious dependencies between variables that are marginally independent. Such associations provide a clear empirical example of the well-known Berkson's paradox [5], [6]. Our results confirm that collider correction removes these spurious connections, as previously demonstrated in [7]. Figures 2C and 2D show cases in which collider correction or naïve $\rho_{PC}$ generate FN. In fig. 2C the collider correction overcompensates in a tightly coupled structure: the GT network includes a collider node (node 2) between 1 and 3, which are also directly connected. While $\rho_{PC}$ correctly recovers this direct interaction, $\rho_P$ fails to detect it (FN), and the subsequent collider correction, based on agreement between the two, removes the connection. This example highlights the trade-off between sensitivity and specificity introduced by the correction procedure. In Fig. 2D, nodes simultaneously participate in multiple interaction motifs forming a closed loop, and the conditioning set includes nodes belonging to consecutive causal chains. We observe that $\rho_{PC}$, by regressing out the contribution of both nodes 1 and 2, removes variance shared between directly interacting nodes (1 and 3), causing a FN. This illustrates how conditioning on multiple related variables can lead to overcorrection when shared variance reflects true interactions.

The toy examples conceptually simplify why no estimator can be interpreted in purely motif wise terms once multiple interacting structures and incomplete conditioning sets are considered. They also motivate the systematic evaluation of the estimators in high dimensional connectome simulations.

We then evaluated how estimator performance varies as a function of the number of timepoints available for estimation (Fig.3). As the number of timepoints increases, the total number of statistically significant connections also increases. This effect is reflected in both an increase in TPR and a decrease in FNR. However, we also found a slightly increase in FPR, with the largest one occurring for $\rho_P$. Nevertheless, increasing the number of timepoints improves the stability of covariance estimation and resulted in a general increase in balanced accuracy (BA) for all methods except $\rho_P$, which shows a relatively stable trend across sample sizes. We also observed that, in low-sample regimes (up to approximately 600 timepoints), PCA-based regularization provides the most stable behaviour in terms of FPR among the collider-uncorrected estimators and yields the largest improvement in BA compared with standard $\rho_{Pc}$. This is particularly relevant because many applications usually do not allow for long scanning sessions, especially in clinical studies. Beyond this point, BA for $\rho_{PC}$ starts to increase more rapidly, and the performance of the two methods became comparable around 1000 timepoints. Finally, the collider-corrected estimators achieved the best performance in terms of FPR, with values consistently around 0.1.

#### 3.2. High dimensionality networks

Simulation details of high dimensionality networks are reported in section E of the main paper. Following the same implementation, we performed additional analyses using the following parameters combinations:

1) D = 5%, 230 regions, 80% explained variance, no filtering in [0.01,0.1] Hz
2) D = 5%, 345 regions, 99% explained variance, no filtering in [0.01,0.1] Hz
3) D = 5%, 345 regions, 99% explained variance, with filtering in [0.01,0.1] Hz
4) D = 30%, 345 regions, 99% explained variance, no filtering in [0.01,0.1] Hz
5) D = 30%, 345 regions, 99% explained variance, with filtering in [0.01,0.1] Hz

We aggregated the estimated connectivity matrices across the 25 repetitions and derived a group connectome for $\rho_P$, $\rho_{PC}$, $PCA - \rho_{PC}$, as for the experimental fMRI dataset. Here, we show the results in radar plots (Figure 4) reporting precision, sensitivity, specificity, FPR, FNR, F1 score, Matthew's correlation coefficient (MCC), and BA.

##### 3.2.1. Low-collinearity scenario

In the first scenario (DAG density 5%, 230 regions, 80% PCA variance retained, no filtering), collinearity among timeseries was minimal. In this setting, standard $\rho_{Pc}$ shows slightly better performance than $PCA - \rho_{Pc}$, with a similar pattern observed for their collider-corrected counterparts. Pearson correlation consistently exhibits the poorest performance across metrics.

##### 3.2.2. Increasing number of regions and explained variance in PCA

In the second scenario (DAG density 5%, 345 regions, 99% PCA variance retained, no filtering), increasing the amount of variance retained in the PCA-based conditioning set improves the performance of $PCA - \rho_{Pc}$. Specifically, it achieves performance comparable to standard $\rho_{PC}$ across the evaluated metrics, while $\rho_P$ continues to show inferior results.

##### 3.2.3. Influence of filtering

We then evaluated the effect of temporal filtering (0.01–0.1 Hz). In this scenario (DAG density 5%, 345 regions, 99% PCA variance retained, with filtering), the overall behaviour remaines consistent with the previous condition, but all estimators show slightly improved absolute metric values.

##### 3.2.4. Collinearity induced by network structure

When collinearity is imposed by the weights of the GT network (DAG density 30%, 345 regions, 99% PCA variance retained, no filtering), $PCA - \rho_{PC}$ outperformes standard $\rho_{PC}$ across most of the evaluated metrics.

##### 3.2.5. Combined effects of collinearity and filtering

Finally, we evaluated a scenario combining high network density and temporal filtering (DAG density 30%, 345 regions, 99% PCA variance retained, with filtering). In this case, $PCA - \rho_{PC}$ continues to show improved performance relative to $\rho_{PC}$, although the differences between the methods are smaller compared to the previous condition.

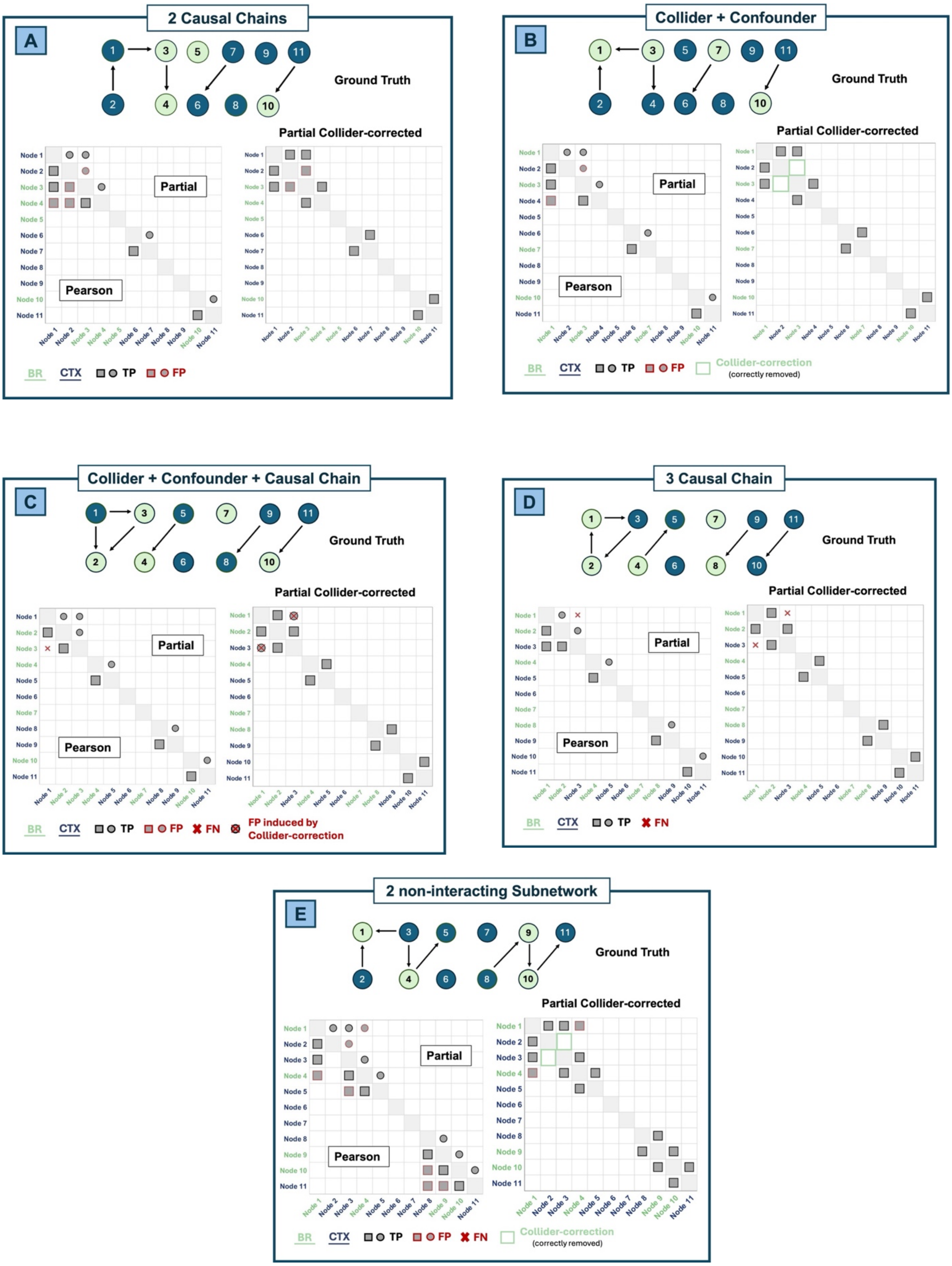


*Supplementary Figure 2: Toy examples illustrating the behaviour of Pearson, partial, and collider-corrected partial correlation in multiple causal motifs. Top panels show the ground truth (GT) directed networks; bottom panels report the estimated connectivity matrices. Color-coded markers indicate correctly identified connections (true positive), spurious connections (false positive), missed connections (false negative), connections correctly or incorrectly removed by collider correction, according to the legend shows in the bottom part of each subfigure. Cortex nodes labels are shown in blue (CTX), whereas brainstem-like nodes are reported with green labels (BR). A) Network with 2 causal chains; B) Network with collider and confounder; C) Network with collider, confounder and causal chain; D) Network with 3 causal chains; E) Network with 2 subnetworks without any interaction between them.*

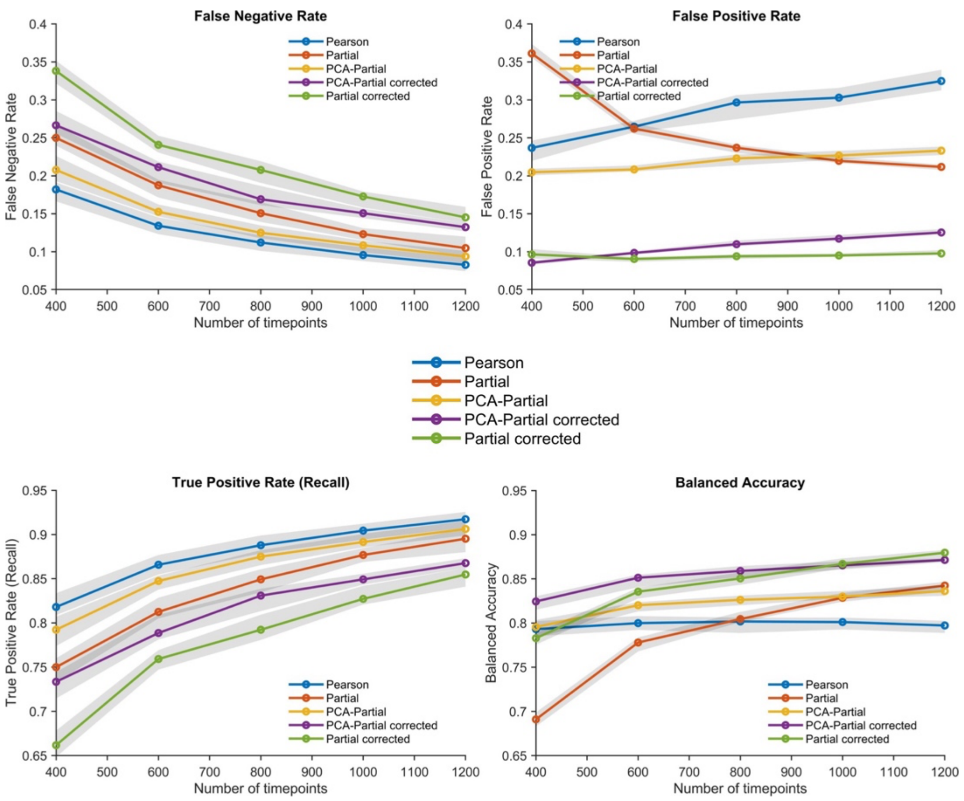


*Supplementary Figure 3: Effect of the number of timepoints. Performance of FC estimators used in this work as a function of the number of timepoints used to generate timeseries. The panels report false negative rate (FNR), false positive rate (FPR), true positive rate (TPR/recall), and balanced accuracy (BA). Solid lines indicate median values across simulations, and shaded areas represent interquartile ranges.*

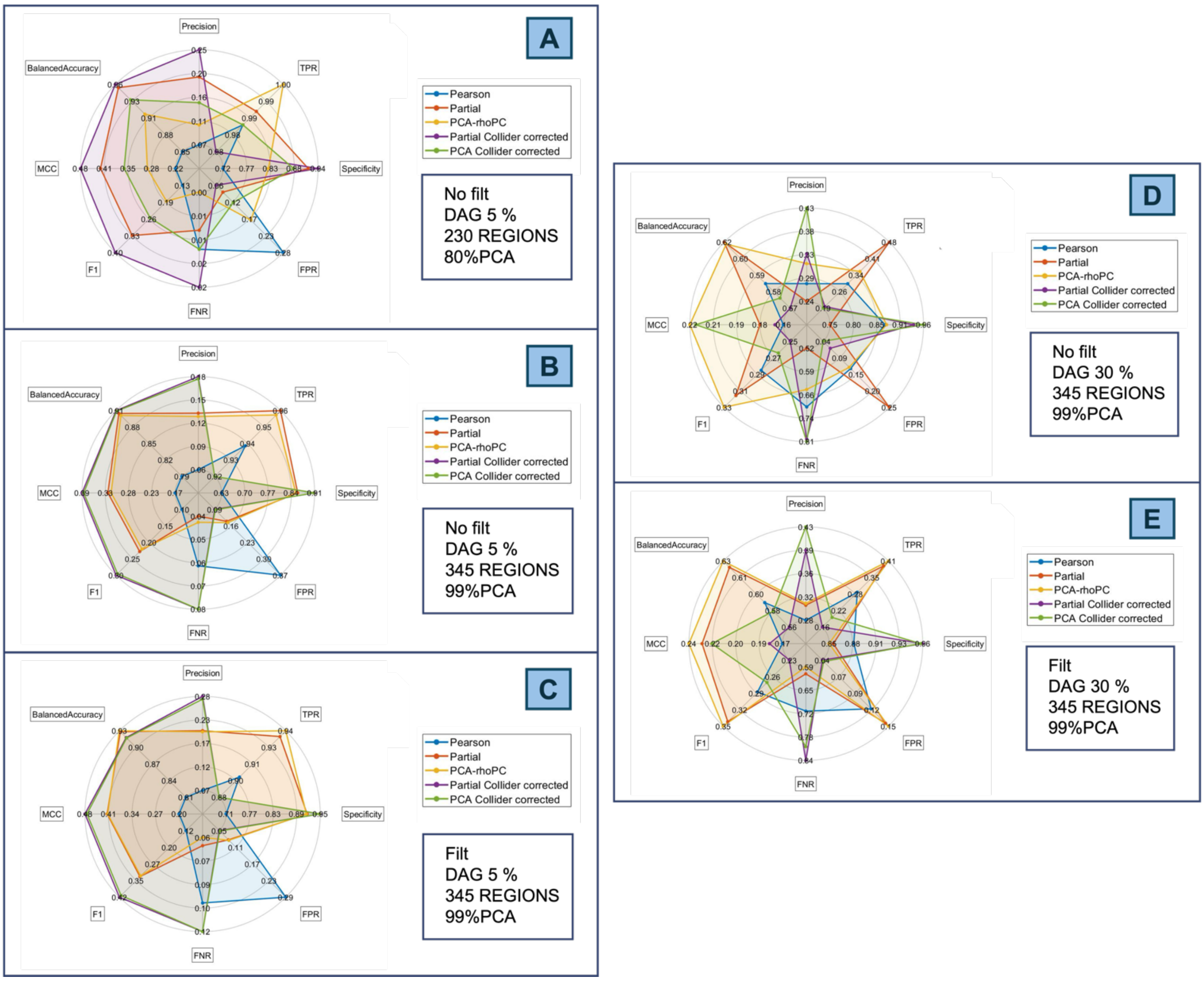


*Supplementary Figure 4: Radar plots summarizing the performance of different FC estimators used in this work across representative simulation conditions. Each axis of the radar plot represents a performance metric including precision, sensitivity, specificity, FPR, FNR, F1 score, MCC and BA. A) low density network (DAG 5%), 230 regions, 80% PCA variance retained, no filtering; B) low density network (DAG 5%), 345 regions, 99% PCA variance retained, no filtering; C) low density network (DAG 5%), 345 regions, 99% PCA variance retained, with temporal filtering (0.01-0.1 Hz); D) high density network (DAG 30%), 345 regions, 99% PCA variance retained, no filtering; E) high density network (DAG 30%), 345 regions, 99% PCA variance retained, with temporal filtering (0.01-0.1 Hz).*

### 4. EXPERIMENTAL 3T fMRI DATASET

#### Quality control and subject exclusion

Quality control (QC) was performed at the run level using motion- and signal-based metrics computed from the EPI timeseries, together with the residual model degrees of freedom (DoF) after preprocessing. As part of the original data release, runs had already undergone an initial motion-based QC, and subjects were retained only if mean frame-to-frame displacement was ≤ 0.15 mm [8]. We then applied additional run-level QC more tailored to brainstem FC. For each run, we required: (*i*) limited global signal fluctuations, quantified as the proportion of frames exceeding a data-driven sDVARS threshold computed on the scaled EPI timeseries, with runs excluded when the fraction of suprathreshold frames was >5%; and (*ii*) adequate temporal signal-to-noise ratio (tSNR) within a cortical mask and a subcortical mask (brainstem, diencephalon, and basal ganglia regions), computed both on minimally processed EPI data (to flag acquisition-related signal dropout) and on the final timeseries (to assess residual signal stability). For both masks, run-wise tSNR was required to exceed the 5th percentile of the corresponding sample distribution. Because nuisance regression was performed jointly with band-pass filtering and timepoint censoring, we also evaluated the residual DoF after regression to confirm that the model was not over-parameterized and that non-zero residual DoF remained for subsequent connectivity estimation (see [9] and afni_proc documentation for more details). Subject-level exclusion was performed by removing participants for whom at least one of the two runs failed any QC criterion, ensuring that all retained subjects contributed two runs of comparable quality for subsequent brainstem FC analyses. After QC, 16 participants (two runs each) were retained for further analysis.
Figure 5 shows tSNR maps from a representative subject (panel A) and pooled tSNR distribution across subjects and runs (panel B). tSNR was evaluated within a whole-brain mask and within an ad hoc subcortical mask including brainstem regions.

Across subjects, tSNR increased markedly after nuisance regression compared with minimally preprocessed data. As expected for 3T acquisitions, cortical tSNR was higher than subcortical tSNR [10][11]; however subcortical tSNR values remained within a range reported in prior brainstem fMRI studies and compatible with reliable ROI-based connectivity estimation [12], [13]. Across all subjects and runs, nuisance regression substantially increased tSNR in both cortical and subcortical regions [11] with only three runs falling below the 5th-percentile threshold.

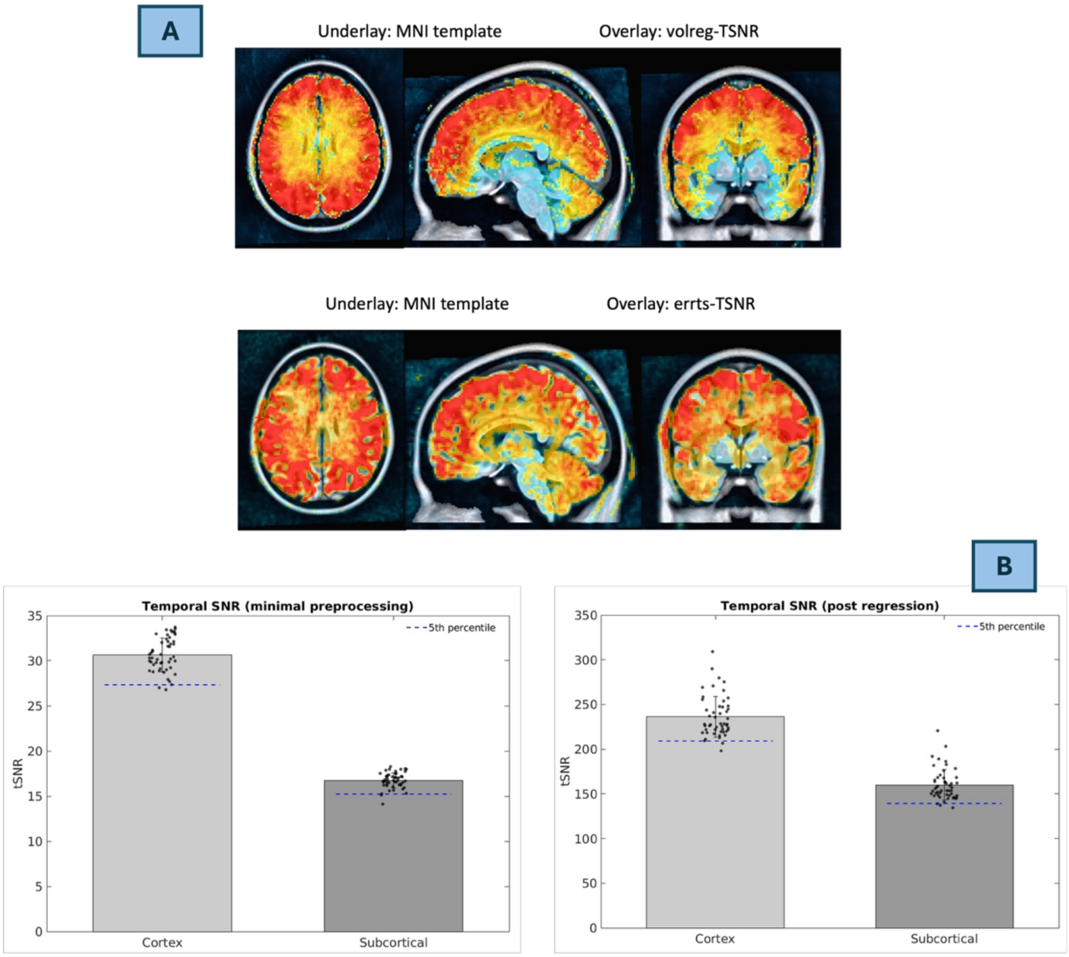


*Supplementary Figure 5: Temporal SNR before and after nuisance regression.A) Underlay: MNI template. Overlay: tSNR estimated from the motion-corrected BOLD time series (top, minimally pre-processed data) and from the residual time series after nuisance regression (bottom). Axial, sagittal, and coronal views from a representative subject are shown, illustrating full brainstem coverage and the substantial increase in tSNR after regression. B) Group summary of tSNR in cortical and subcortical masks (including brainstem regions). Bars show mean tSNR across subjects and runs for cortex and subcortical regions before (left) and after (right) nuisance regression. Dots represent individual run values. The dashed blue line indicates the 5th percentile of the corresponding distribution in each panel.*

**Brainstem nuclei used in the connectomic analysis**

| LABEL | REGION NAME |
|---|---|
| BRAINSTEM NUCLEI | |
| **Cli_RLi** | **Caudal-rostral linear raphe** |
| CnF | Cuneiform nucleus |
| **DR** | **Dorsal raphe** |
| IC | Inferior Colliculus |
| iMRt | Inferior medullary reticular formation |
| ION | Inferior olivary nucleus |
| isRt | Isthmic reticular formation |
| LC | Locus coeruleus |
| LDTg_CGPn | Laterodorsal tegmental nucleus – central gray of the rhomboencephalon |

| | |
|---|---|
| LPB | Lateral parabrachial nucleus |
| MiTg_PBG | Microcellular tegmental nucleus - prabigeminal nucleus |
| **MnR** | **Median raphe** |
| MPB | Medial parabrachial nucleus |
| mRt | Mesencephalic reticular formation |
| **PAG** | **Periaqueductal gray** |
| PCRtA | Parvicellular reticular nucleus Alpha part |
| **PMnR** | **Paramedian nucleus** |
| PnO_PnC | Pontine reticular nucleus, oral and caudal parts (pontis oralis and caudalis) |
| PTg | Pedunculotegmental nucleus |
| **RMg** | **Raphe magnus** |
| RN | Red Nucleus |
| RN1 | Red Nucleus: subregion 1 |
| RN2 | Red Nucleus: subregion 2 |
| **ROb** | **Raphe obscurus** |
| **RPa** | **Raphe pallidus** |
| SC | Superior colliculus |
| sMRt | Superior medullary reticular formation |
| SN | Substantia nigra |
| SN1 | Substantia nigra: subregion 1 (reticulata) |
| SN2 | Substantia nigra: subregion 2 (compacta) |
| SOC | Superior olivary complex |
| SubC | Subcoeruleus |
| Ve | Vestibular nuclei complex |
| VSM | Viscero-sensory-motor nuclei complex |
| VTA_PBP | Ventral tegmental area – parabrachial pigmented nucleus complex |

*Table 1: Brainstem ROIs used in this study. List of seed and target regions included in the connectomic analysis. Bold labels refer to midline brainstem nuclei, whereas the remaining regions are bilateral structures and were treated during the analysis as separate left- and right-hemisphere ROIs.*

**Multicollinearity and run-to-run stability assessment**

To characterise run-dependent effects on conditioning, we repeated the multicollinearity analysis reported in the main manuscript separately for run 1 and run 2. For each subject and method, we computed the condition number ($n$) of the standardized regressor covariance matrix. As described in Method section, for $PCA - \rho PC$, $n$ was computed in the PCA-transformed regressor space after dimensionality reduction; for Ridge, $n$ was computed for the penalized system; for Lasso and Elastic Net, $n$ was computed on the active set of predictors (non-zero coefficients). Across-subject summaries (median, IQR, and MAD) are reported in Tables S2-S3.

Multicollinearity among covariate timeseries inflates the variance of regression coefficients [14] and can therefore propagate to partial correlation estimates. Given the direct relation between the β regression parameters and partial correlation coefficients ($\rho_{PC}\left(x_i, x_j \middle| x_{-(i,j)}\right) = \frac{\beta_{ij} SD(\epsilon_j)}{SD(\epsilon_i)}$) [15], this effect may manifest as increased between-subject dispersion of estimated FC. We therefore summarized the group mean and between-subject standard deviation of edge weights for each method ($\rho_p$, standard $\rho_{PC}$, $PCA - \rho_{PC}$, $\rho_{L2}$, $\rho_{L1}$ and $\rho_{EN}$), restricted to brainstem connections (brainstem–to-brainstem upper triangle and brainstem–to-cortex edges). These summaries characterise how estimator choice affects the overall scale and dispersion of connectivity weights across subjects, and reflect a mixture of intrinsic inter-individual variability, noise, and estimator instability (Figure 6). We observed that standard partial correlation shows a larger between-subject dispersion than PCA-based partial correlation, L2-partial correlation and Pearson correlation. In the single run analysis, Lasso and Elastic Net show STD comparable to naïve partial correlation. Moreover, in run 1, the distribution of mean connectivity values is broader for standard partial correlation and sparse regularization than for the other estimators; this difference is attenuated when runs are pooled.

Finally, we quantified within-subject stability. We used the same pipeline of the aggregated runs to estimate connectomes separately from run 1 and run 2. For each method, we selected the set of top-K edges (here K=500) based on the absolute group mean Fisher-z. For each subject, on the selected edges, we compared the connectomes from the two runs using two complementary metrics: *(i)* brainstem pattern-level reliability, quantified as the Pearson correlation between the two run specific edge weight vectors, and *(ii)* brainstem run-to-run absolute change, defined as the median absolute difference across the selected edges ($\Delta_s = median_{e \in TopK} |z_e^{run1} - z_e^{run2}|$). Both metrics were applied on the upper triangle of brainstem-to-brainstem connectivity and the brainstem-to-cortex edges.

Figure 7A shows pattern-level reliability results and figure 7B instead reports edgewise run-to-run absolute change. Pearson correlation shows high pattern-level reproducibility (median=0.771; IQR=0.081). $PCA - \rho_{PC}$ improvs run-to-run reliability compared to standard $\rho_{PC}$ (median=0.697; IQR=0.158 vs median=0.416; IQR=0.227), indicating that PCA regularization increases the stability of multivariate conditioning. Ridge achieves reliability closest to Pearson (median=0.772; IQR=0.081), whereas sparse penalties exhibits lower pattern-level reproducibility (Lasso: median=0.377; IQR=0.245; Elastic Net: median=0.407; IQR=0.243).

A similar ranking emerges for edgewise stability. Pearson and Ridge yield the smallest run-to-run changes (median $\Delta_s$= 0.1533 and 0.1534, respectively). $PCA - \rho_{PC}$ shows intermediate changes (median $\Delta_s$= 0.1776), while $\rho_{PC}$, Lasso and Elastic Net show larger run-to-run differences (median $\Delta_s$= 0.2808; 0.3253; 0.3459, respectively). For completeness, mean ± SD summaries of $\Delta_s$ and additional diagnostic analyses are provided in Tables S4–S5.

For standard $\rho_{PC}$ (and, to a lesser extent, sparse regularization), the discrepancy between median-based and mean-based summaries suggests that a limited number of extreme run-to-run edge-weight differences inflate mean ± SD, whereas the median better captures typical subject-level behaviour. As a diagnostic for edge-weight saturation potentially linked to numerical instability, we quantified the fraction of Top-K edges with near-perfect connectivity estimates (|r|>0.99) in either run (Table S5). Specifically, we report: (i) the overall fraction pooling all Top-K edges across all subjects, (ii) the percentage of subjects showing at least one near-perfect edge, and (iii) the maximum subject wise fraction (worst case). This fraction is negligible for Pearson, $PCA - \rho_{PC}$, and Ridge (overall fractions on the order of $10^{-3}$), but is higher for the other estimators. In particular, the worst-case subject-wise fraction is markedly larger for standard $\rho_{PC}$ (max = 0.7740) than for other methods ($PCA - \rho_{PC}$ max = 0.0160; Pearson and Ridge max = 0.0020). These results provide an interpretable explanation for the discrepancy between median-based and mean-based run-to-run summaries: methods prone to extreme edge weights, consistent with ill-conditioned estimation arising from multicollinearity, yield skewed distributions of the run-to-run variability and inflated mean ± SD.

| | ***Partial Correlation*** | ***PCA Regularization*** | ***L1 Penalty*** | ***L2 Penalty*** | ***EN Penalty*** |
|---|---|---|---|---|---|
| Median ± IQR | $2.88e^3 \pm 3.85e^3$ | 12.27 ± 4.38 | $1.46e^3 \pm 2.73e^3$ | $1.16e^3 \pm 1.96e^3$ | $1.49e^3 \pm 2.81e^3$ |
| MAD | $3.72e^{12}$ | 4.00 | $2.55e^{12}$ | $2.09e^{12}$ | $2.66e^{12}$ |

*Table S2: Multicollinearity summary for run 1*

| | ***Partial Correlation*** | ***PCA Regularization*** | ***L1 Penalty*** | ***L2 Penalty*** | ***EN Penalty*** |
|---|---|---|---|---|---|
| Median ± IQR | $5.24e^3 \pm 1.87e^{13}$ | 13.08 ± 4.30 | $2.93e^3 \pm 1.26e^{13}$ | $2.22e^3 \pm 1.31e^{13}$ | $2.99e^3 \pm 1.44e^{13}$ |
| MAD | $1.05e^{13}$ | 2.36 | $6.82e^{12}$ | $7.92e^{12}$ | $7.66e^{12}$ |

*Table S3: Multicollinearity summary for run 2*

**A**

**Pooled across runs**

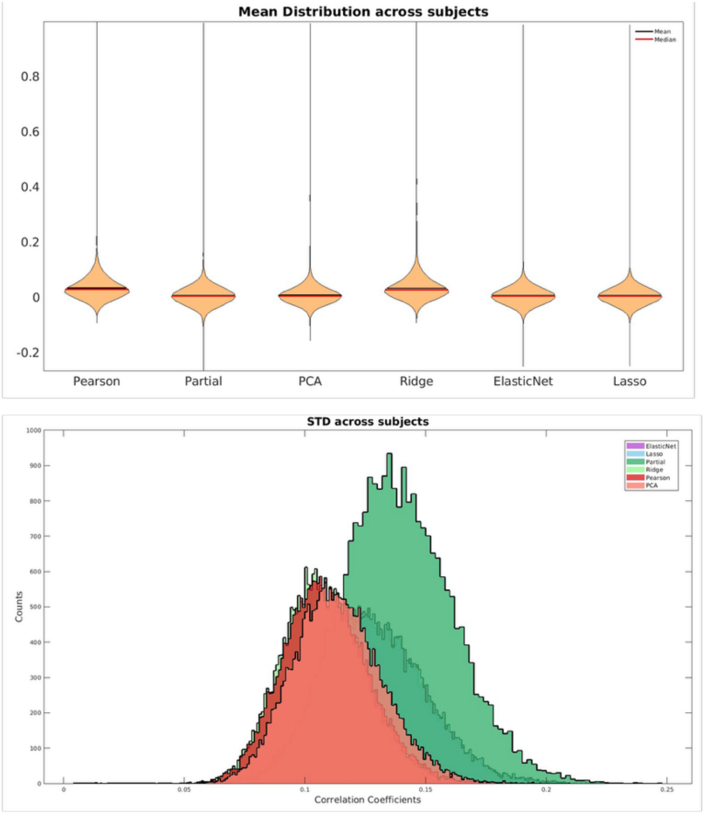


**B**

**Run 1**

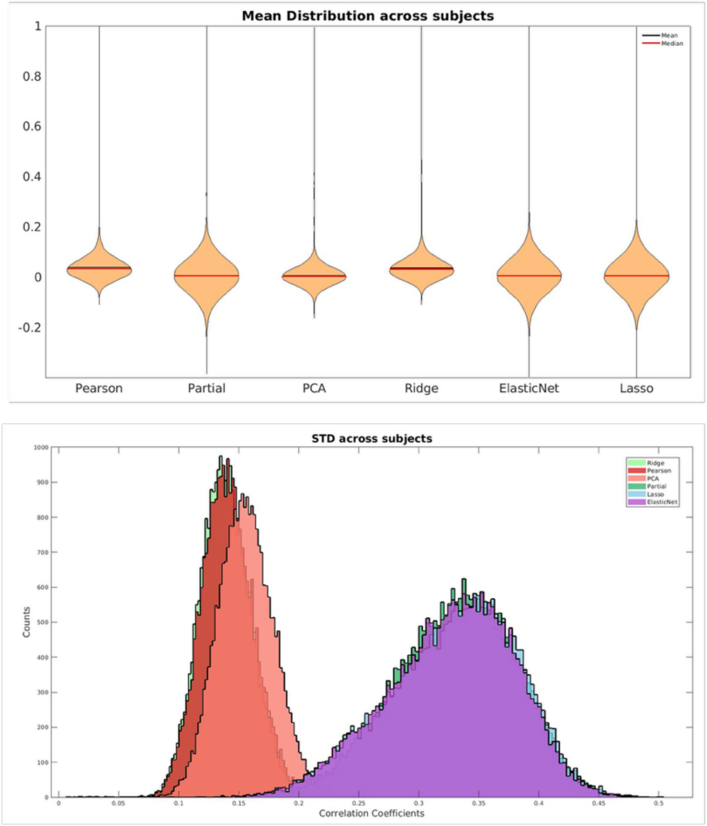


*Supplementary Figure 6: Average values and standard deviation of edge weights for Pearson correlation, standard partial correlation, PCA-based partial correlation, and regularized partial correlation (Ridge, Lasso, Elastic Net), restricted to brainstem–brainstem (upper triangle) and brainstem–cortex edges. Mean and SD were computed across subjects on these brainstem connectivity estimates. A) Results obtained on the FC estimation from data concatenation of run 1 and run 2; B) Results for a single run.*

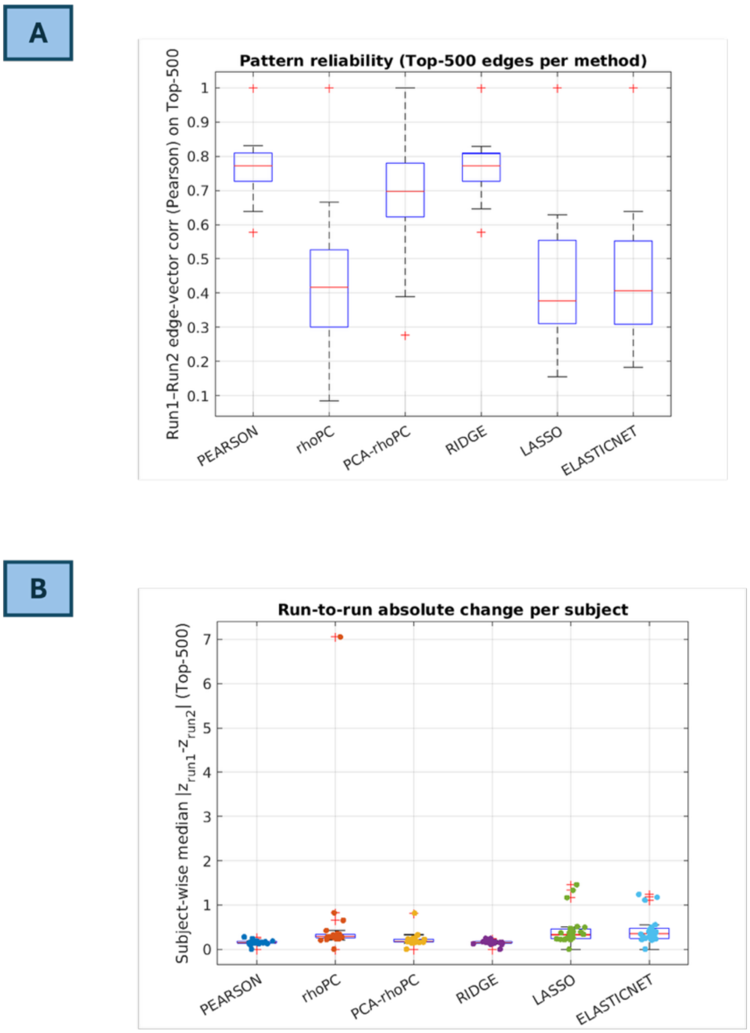


*Supplementary Figure 7: Within-subject run-to-run stability of the brainstem connectome across estimators. A) Pattern-level reliability, computed as the Pearson correlation between run-specific Top-K edge-weight vectors (K = 500). B) Edgewise run-to-run change, quantified as the subject-wise median across Top-K edges of $|z_{run1} - z_{run2}|$, where z denotes Fisher-z-transformed edge weights. For each method, Top-K edges were defined using the absolute group-mean Fisher-z (computed across both runs). Boxplots summarise distributions across subjects.*

| ***Method*** | ***Median ± IQR*** | ***Mean ± SD*** | ***Skew Indicator*** |
|---|---|---|---|
| Pearson Corr | 0.153 ± 0.039 | 0.155 ± 0.050 | 1.01 |
| Partial Corr | 0.281 ± 0.081 | 0.592 ± 1.385 | 2.11 |
| PCA Partial Corr | 0.178 ± 0.068 | 0.209 ± 0.143 | 1.18 |
| Ridge Corr | 0.154 ± 0.036 | 0.153 ± 0.046 | 1.00 |
| Lasso Corr | 0.325 ± 0.222 | 0.436 ± 0.362 | 1.34 |
| Elastic Net Corr | 0.346 ± 0.227 | 0.424 ± 0.313 | 1.23 |

*Table S4: Run-to-run variability ($\Delta_s$) across subjects for Top-K edges. Distribution of $\Delta_s$ (K = 500) are summarised using median ±IQR and mean ±SD across subjects. The skew indicator is defined as Mean/Median.*

| ***Method*** | ***Overall fraction*** | ***Subject with ≥ 1(%)*** | ***Subject max*** |
|---|---|---|---|
| Pearson Corr | 0.0008 | 37.5 | 0.0020 |
| Partial Corr | 0.0415 | 33.3 | 0.7740 |
| PCA Partial Corr | 0.0010 | 20.8 | 0.0160 |
| Ridge Corr | 0.0007 | 33.3 | 0.0020 |
| Lasso Corr | 0.0236 | 29.2 | 0.2180 |
| Elastic Net Corr | 0.0173 | 41.7 | 0.1520 |

*Table S5: Diagnostic of near-perfect edge weights. For each method, we quantified the prevalence of near-perfect connectivity estimates within the Top-K edges (K=500; selected per method based on the absolute group-mean Fisher-z). An edge was classified as near perfect if |r|>0.99 in either run. "Overall fraction" reports the fraction of near-perfect entries pooling all Top-K edges across all subjects. Subject-wise fractions $f_{SUBJ}$ were computed as the proportion of Top-K edges classified as near-perfect for each subject; we report "Subjects with ≥1", the percentage of subjects with at least one near-perfect edge ($f_{SUBJ} > 0$), and "Subject max", the maximum $f_{SUBJ}$ across subjects.*

**Connectome of the Inferior Colliculus with Pearson and PCA-based Partial Correlation**

Following the scheme of the main manuscript, we here report the functional cortical and subcortical connectomes for Inferior Colliculus (IC) estimated using both Pearson and PCA-based partial correlation. Supplementary Figure 8 shows that for $\rho_P$, connectivity extends beyond temporal regions to include occipital, parietal, and hippocampal regions. $PCA - \rho_{PC}$ reduces visual contributions and instead more selectively engages premotor regions, somatosensory and sensorimotor cortex, inferior parietal cortex, and medial temporal regions (pre/parasubiculum and parahippocampal areas). However, an absence of extended auditory cortical coupling is observed. Within brainstem FC, IC shows significant interactions with CnF, DR, PAG, MITg/PBG and RN. IC is classically described as a key midbrain relay of the ascending auditory pathway, projecting to the medial geniculate nucleus (MG) and subsequently to auditory cortex [16]. Consistent with this organization, $\rho_P$ revealed IC–MG coupling and widespread connectivity with temporal and multimodal cortical regions. However, this broad connectivity profile likely reflects shared network fluctuations rather than exclusively direct interactions. After controlling for multicollinearity, $PCA - \rho_{PC}$ revealed a markedly more selective and interpretable network organization. Specifically, IC connectivity was centred on sensorimotor and opercular regions, including primary somatosensory cortices, posterior parietal areas (areas 1–3, 5L/5m), premotor cortex (area 6), and secondary somatosensory/opercular regions (SII, OP areas). This pattern is consistent with evidence that the IC is not a purely feedforward auditory relay, but a multisensory hub involved in the integration of auditory information with somatosensory, motor and orienting signals [17] Importantly, IC–MG connectivity observed with $\rho_P$ was not retained in the $PCA - \rho_{PC}$ connectome. This finding may indicate that IC–thalamic coupling, while present and expressed at the level of shared covariance, does not persist as a strong residual dependency after conditioning (i.e., once the contribution of the broader brainstem network is controlled). This observation aligns with the known organization of the auditory thalamus, which is embedded in a complex circuit involving multiple inputs, including cortical feedback and non-lemniscal pathways [18], [19]. Interestingly, $PCA - \rho_{PC}$ retained connectivity with parahippocampal and presubicular regions, suggesting a potential link between IC activity and contextual or spatial processing systems. Although likely indirect, this may reflect integration of auditory signals with higher order representations of context and environment [20], [21]. Finally, the absence of strong temporal auditory connectivity in the $PCA - \rho_{PC}$ connectome may reflect methodological factors: temporal lobe regions are known to be affected by susceptibility artifacts in fMRI, potentially reducing sensitivity.

Overall, these findings indicate that Pearson correlation captures the IC within a broad multimodal covariance structure, whereas $PCA - \rho_{PC}$ isolates a subset of interactions revealing a functional organization in which the IC also participates in distributed sensorimotor and multisensory networks.

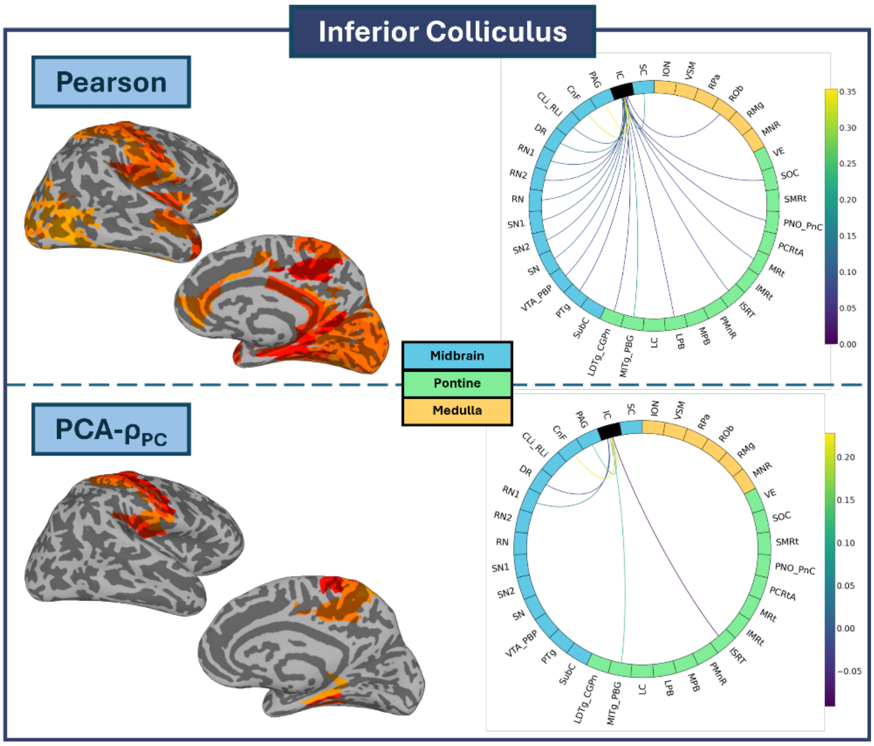


*Supplementary Figure 8: Functional connectome of the Inferior Colliculus obtained using Pearson and PCA-based partial correlation. A) Statistically significant cortical connections projected onto the cortical surface; B) circular plots of significant connections between each nucleus and other brainstem nuclei and subcortical areas.*

### Ridge Functional Connectomes

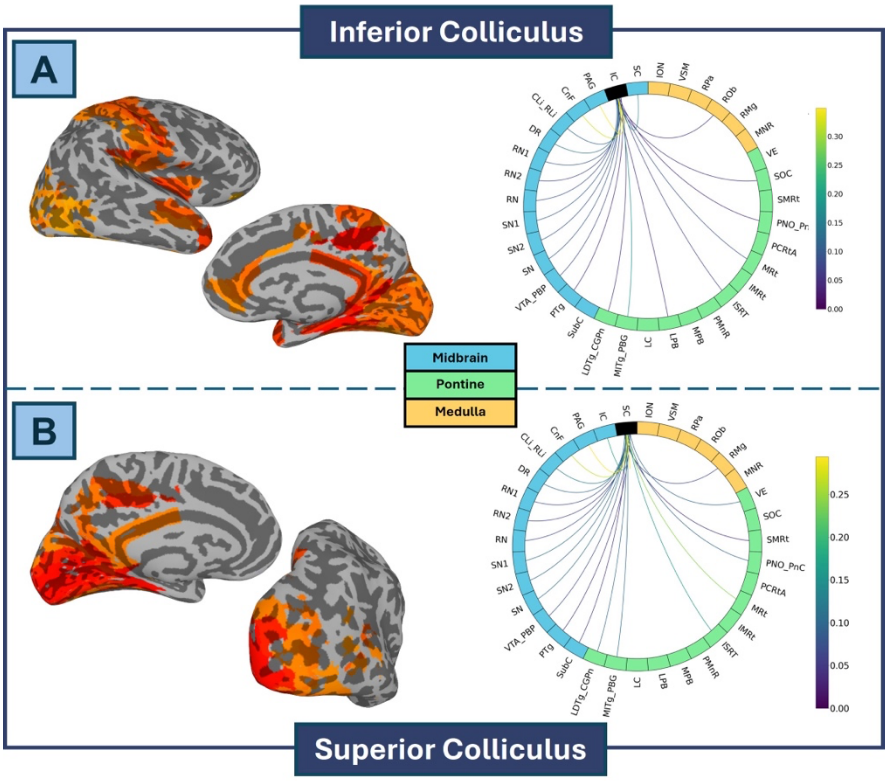


*Supplementary Figure 9: Functional connectomes obtained using partial correlation and Ridge regularization for A) Inferior Colliculus (IC); B) Superior Colliculus (SC). In each panel, the upper section shows statistically significant cortical connections projected onto the cortical surface, while the lower section displays circular plots of significant connections between each nucleus and other brainstem nuclei and subcortical areas.*
